# ASCH Domain-Containing Proteins Act as tRNA *N*^4^-acetylcytidine Erasers

**DOI:** 10.64898/2026.02.27.708502

**Authors:** Roberta Statkevičiūtė, Mikas Sadauskas, Agota Aučynaitė, Audrius Laurynėnas, Greta Gakaitė, Rolandas Meškys

## Abstract

*N*^4^-acetylcytidine (ac^4^C) is a conserved RNA modification that enhances RNA stability and translation accuracy. Emerging evidence suggests that ac^4^C levels can change in response to cellular and environmental triggers. Despite these indications of regulatory dynamics, the enzymes responsible for removing ac^4^C, beyond the recently proposed rRNA deacetylase SIRT7, remain largely unknown. Here, we examined 19 ASCH domain-containing proteins from bacteria, archaea, and humans to determine their possible activity in tRNA deacetylation. Despite differences in their sequences, structures, and nucleic-acid binding properties, all tested proteins were capable of removing ac^4^C from tRNA, revealing a conserved deacetylase activity across diverse species. The proteins were found to vary in nucleic acid recognition, including an archaeal specific helix–turn–helix domain that promotes strong tRNA binding. Together, these findings establish ASCH proteins as a widespread and previously unrecognized family of tRNA deacetylases, suggesting that enzymatic ac^4^C turnover may require complex regulation within the cells.

**GRAPHICAL ABSTRACT:** 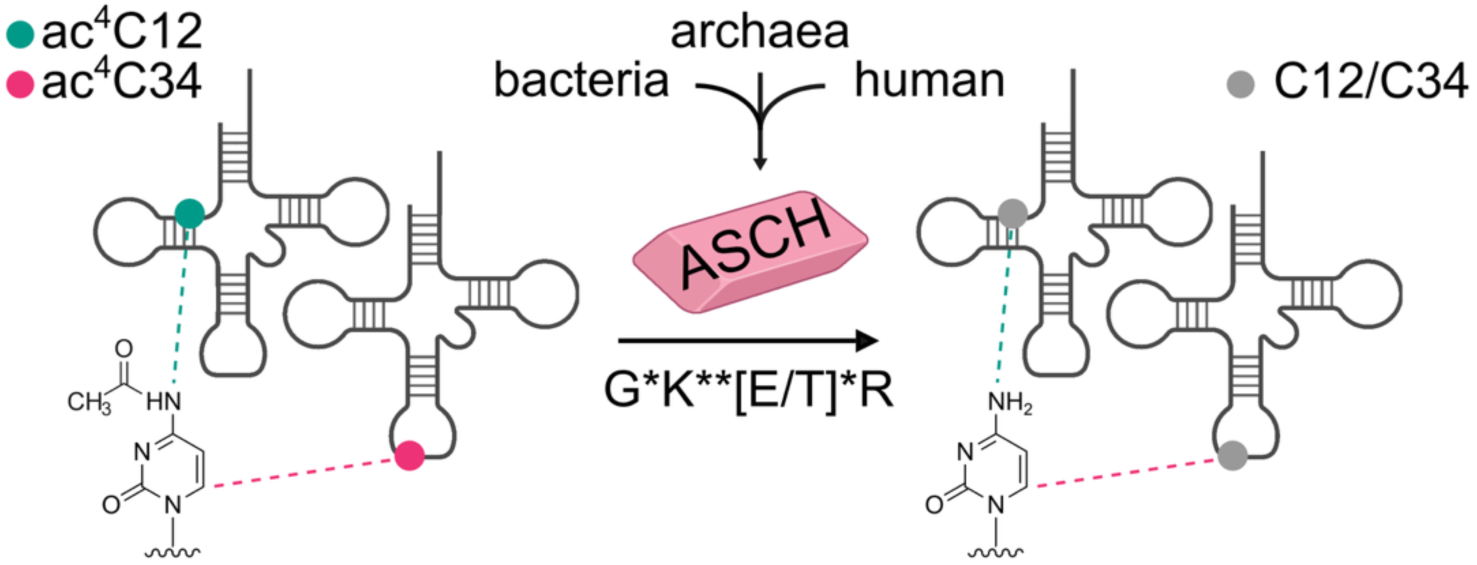

## INTRODUCTION

Post-transcriptional modifications of coding and non-coding RNA molecules are crucial for their processing, structural stability, and highly regulated RNA-protein interactions. Recent transcriptomic studies have uncovered more than 170 distinct post-transcriptional modifications across RNA species, of which about 80% have been extensively characterized in transfer RNA (tRNA) (1, 2). Growing evidence suggests that some RNA modifications are dynamic, thus being introduced or removed from RNA molecules as a response to a particular environmental stressor (temperature, oxidative stress, nutrient deficiency), signaling that these chemical alterations could play a part in rapid stress-induced cellular adaptation. Although the biosynthetic pathways of many modified nucleotides are well studied, the catabolism of these compounds is less understood. Moreover, most identified modifications are still considered static, as the enzymes involved in their removal (erasers) have not yet been discovered. Only three RNA de-modification events are known today: demethylation of *N*^1^-methyladenosine (m^1^A) and *N*^6^-methyladenosine (m^6^A), and desulfidation of 4-thiouridine (s4U). In humans, the demethylation of m^1^A has been shown to be catalyzed by the α-ketoglutarate-dependent demethylases ALKBH1 and ALKBH3 (3, 4), while m^6^A removal is catalyzed by the demethylases FTO and ALKBH5 (5, 6). Recently, the first bacterial tRNA de-modifying enzymes emerged. Hence, the m¹A demethylases RMD1 and RMD2, involved in stress responses, and the s4U desulfidase RudS, participating in the UV-induced damage response, have been identified and characterized (7, 8).

*N*^4^-acetylcytidine (ac^4^C) is a conserved post-transcriptional RNA modification found across all domains of life. In eukaryotes, ac^4^C is present in 18S ribosomal RNA (rRNA) (9), the 12^th^ position of the D-arm of tRNA^Leu/Ser^ (10–13), and there are emerging studies providing evidence of acetylated positions in messenger RNAs (mRNA) (14–16). The *N*-acetyltransferase NAT10 in humans, together with its homologs in other eukaryotes, such as Kre33 in yeasts, catalyzes ac^4^C formation with the help of adapter proteins or snoRNAs (17, 18). It has been demonstrated that ac^4^C is essential for eukaryotic ribosome biogenesis and transcript stability, particularly during heat shock, as it facilitates Watson-Crick pairing with guanosine in double-stranded RNA motifs, thereby enhancing the translation process (19). In mRNA, the regulatory effect of ac^4^C is location-specific: in the 5’ untranslated regions (5’-UTRs), it can form translation-inhibiting structures and inhibit start codons (20); however, when the modification is within the coding sequences (CDSs), it enhances translation efficiency by providing stability and, therefore, prolonging the half-life of the transcripts (17). It has been shown that the acetylation of host or viral transcripts can contribute to the progression of viral infection by promoting virus replication in host cells (22, 23). Downregulation of NAT10 reduces the severity of infection. Additionally, due to RNA degradation in an oxidative stress environment, ac^4^C can act as a signaling molecule in pro-inflammatory processes and activate the NLRC4 and NLRP3 inflammasomes (24–26). In humans, increased levels of NAT10 lead to an increase of ac^4^C in serum and urine as a potential biomarker for diabetes and certain types of cancer (27, 28). This suggests that NAT10/ac^4^C could be potential therapeutic targets; however, while much is known about the synthesis of ac^4^C, the catabolism pathways remain mostly unknown. Recent research on the HEK293T cell line has indicated that SIRT7, a member of the sirtuin family of histone deacetylases, may act as an eraser for rRNA ac^4^C, offering new insights into the regulation mechanisms of this modification (29).

In bacteria, ac^4^C is primarily associated with the anticodon region of tRNA, where it was first identified at 34^th^ position of methionine elongator tRNA (tRNA^eMet^) (30). In γ-proteobacteria, the NAT10 homolog TmcA modifies this wobble position, therefore ensuring translation accuracy by strengthening C-G pairing between the codon and anticodon (31). In some bacteria, including *Bacillus subtilis*, that lack the *tmcA* gene, an acetate-dependent acetylation is catalyzed by aminoacyl-tRNA synthetase TmcAL (32). Homologs of TmcA have also been identified in archaea (33), where ac^4^C modifications are introduced at multiple sites within tRNA and rRNA (33–36). Although ac^4^C is generally viewed as a stable and permanent RNA modification, growing evidence suggests that its formation is dynamic and can be influenced by environmental conditions. In hyperthermophilic archaea, ac^4^C modifications become more abundant at higher growth temperatures, and deletion of the ac^4^C-synthesizing enzyme causes a temperature-sensitive phenotype with elevated translation errors under heat stress (34, 35, 37)

The *Escherichia coli* amidohydrolase YqfB and its mesophilic homologs from *Buttiauxella agrestis*, *Cronobacter universalis*, *Klebsiella pneumoniae*, and *Shewanella loihica* have previously been shown to hydrolyze the *N*^4^-acylated cytosine and cytidine compounds (38, 39). Based on the kinetic parameters, ac^4^C has been suggested as the primary substrate. It was suggested that amidohydrolytic activity towards *N*-substituted cytidine compounds could be dependent on the catalytic triad composed of Lys21 (general base), Thr24 (nucleophile), and Glu74 (general acid), while Arg26 was suggested to form an atypical oxyanion hole. However, a recently updated crystal structure of *E. coli* YqfB bound to ac^4^C clarified the catalytic mechanism, in which water acts as a nucleophile while Thr24 is involved in its activation (40). These small monomeric amidohydrolases, containing ∼110 amino acids, belong to the human activating signal co-integrator homology (ASCH) superfamily, which may be involved in transcriptional co-activation and RNA regulation. Like ac^4^C, ASCH domain-containing proteins are widespread in nature, but their potential biological functions remain under debate (41). The ASCH domain was first identified as a C-terminal domain of the thyroid hormone receptor interacting protein 4 (ASC-1/TRIP4), which is a component of the multimeric activating signal co-integrator 1 complex (ASCC) involved in the activation of a variety of transcription factors, including serum response factor (SRF), nuclear factor κB (NF-κB), and nuclear hormone receptors (41–44). Although it has recently been demonstrated that the C-terminal ASCH domain of TRIP4 can bind nucleic acids, suggesting a role in transcriptional regulation, ribosome quality control, or DNA damage repair, the cellular function of the domain remains unknown (45). The ASCH domains were also found in human endothelial-overexpressed LPS-associated factors 1 and 2 (EOLA1 and EOLA2). EOLA1 was shown to modulate inflammatory responses by inhibiting LPS-induced interleukin-6 (IL-6) production, thereby controlling excessive inflammation (46, 47). More recently, EOLA1 was reported to be localized in the mitochondrial matrix, where it promotes translation elongation via interactions with 12S mt-rRNA and translation elongation factor TU (48). In contrast, the possible functions of EOLA2 remain hypothetical. In addition, a 147 amino acid-long ZmASCH from *Zymomonas mobilis* was shown to act as a monomeric metal ion-dependent single-stranded RNA ribonuclease, which may impact RNA degradation in the cell (49, 50).

In this study, we characterized a diverse set of 19 ASCH domain-containing proteins originating from human, bacterial, and archaeal sources. Here, we demonstrate for the first time that ASCH domain-containing proteins exhibit a tRNA de-modifying activity targeting ac^4^C modification and uncover an alternative ac^4^C hydrolysis mechanism in the subset of ASCH domain-containing proteins. Furthermore, we identified the presence of a novel helix-turn-helix type nucleic acid-binding domain in archaeal ASCH domain-containing proteins. These findings expand our understanding of the diversity of the ASCH superfamily and highlight tRNA as a common target across phylogenetically distinct ASCH proteins.

## MATERIALS AND METHODS

### Bacterial strains

The *E. coli* DH5α (Novagen, Germany) strain was used for routine DNA manipulation experiments, while the rhamnose-inducible *E. coli* KRX (Promega, USA) strain was used for the synthesis of recombinant target proteins. Parent *E. coli* BW25113 strain, together with its Δ*yqfB* and Δ*tmcA* variants, was obtained from the KEIO Knockout Collection (Japan).

Bacterial strains used in this study for the isolation of the target genes were purchased from the German Collection of Microorganisms and Cell Cultures (DSMZ).

### Growth media

A standard LB broth (Lennox) was used for routine bacteria cultivation and protein synthesis induction (10 g/L tryptone, 5 g/L yeast extract, 5 g/L NaCl). SOB medium (20 g/L Bacto Tryptone, 5 g/L yeast extract, 0.5 g/L NaCl, 0.1 g/L KCl) was used for *E. coli* recovery after transformation.

### Protein expression vectors

The molecular cloning procedures for the genes encoding YqfB and mesophilic YqfB-type amidohydrolases were described in previous publications (38, 39), with the exception of CunASCH, which in this study was truncated by nine N-terminal amino acids to match the UniProt-annotated sequence A0AAC8VS57.

The genes of TthASCH, BmyASCH, and AfuASCH were PCR-amplified using the 2× Phusion Plus PCR Master Mix (Thermo Scientific^TM^, Lithuania, #F631L) and cloned into a C-terminal His-tag containing pLATE31 expression vector by using the aLICator^TM^ LIC Cloning and Expression System Kit 3 (Thermo Scientific^TM^, Lithuania, #K1291). As a source of genomic DNA, intact cells of *Thermus thermophilus* HB8 (DSM 579) and *Bacillus mycoides* (DSM 2048) were used. To amplify the three ASCH domain-containing protein genes from *Archaeoglobus fulgidus* (DSM 4304), purified genomic DNA was used. Primer sequences used in this study are listed in Supplementary Table S1. DNA primers were synthesized by Metabion International AG (Germany) or Azenta Life Sciences (Germany). DNA sequencing was performed by Azenta Life Sciences (Germany).

The genes encoded in the genomes of *Homo sapiens* (EOLA1 isoform 1, TRIP4_ASCH (a stand-alone ASCH domain, residues 435-581), *Pyrococcus kukulkanii* (Pku1-6), and *Zymomonas mobilis* (ZmASCH) were optimized for expression in *E. coli*, chemically synthesized, and cloned into a pET21(+) (TRIP4ASCH, ZmASCH) or pET29b(+) (EOLA1 and Pku1-6) expression vectors by Twist Bioscience (USA). The sequences of expression constructs are listed in Supplementary Table S4.

### Site-directed mutagenesis

Site-directed mutagenesis was performed using a whole-plasmid amplification approach by using 2× Phusion Plus PCR Master Mix (Thermo Scientific^TM^, Lithuania, #F631L) and the respective mutant primers listed in Supplementary Table S2. The resulting PCR products were treated with DpnI FastDigest (Thermo Scientific^TM^, Lithuania, #FD1703) to digest parental DNA, and the products were subsequently purified from agarose gel using the GeneJET Gel Extraction Kit (Thermo ScientificTM, Lithuania, #K0692). The 5′ ends of the purified products were phosphorylated using T4 Polynucleotide Kinase (T4 PNK, Thermo Scientific^TM^, Lithuania, #EK0031). Ligation was performed by self-ligation of the phosphorylated PCR products using T4 DNA ligase (Thermo Scientific^TM^, Lithuania, #EL0011) with overnight incubation at 4°C. The mutant gene variants were sequenced by Azenta Life Sciences (Germany).

### Purification of ASCH domain-containing proteins

A single colony of *E. coli* KRX cells, transformed with the pLATE31 vector harboring a target gene, was inoculated into 5 mL of LB medium supplemented with either ampicillin (50 µg/mL for pET21/pLATE31 constructs) or kanamycin (40 µg/mL for pET29 constructs), and incubated overnight at 37°C with agitation at 180 rpm. Overnight cultures were diluted into fresh, sterile LB medium and grown under the identical conditions until the optical density at 600 nm (OD₆₀₀) reached 0.6–0.8. Protein synthesis was induced by adding 0.5 mM IPTG (Thermo Fisher Scientific, Lithuania, #R0392) and 0.1% L-rhamnose (Sigma-Aldrich, Germany, #W373011), followed by incubation for 16 h at 30°C. Cells were harvested by centrifugation (10 min, 3220 × *g*, 4 °C), resuspended in buffer containing 50 mM Tris-HCl (pH 7.5), 500 mM NaCl, and 5 mM imidazole, and lysed by sonication. The cell debris was removed by centrifugation (20 min, 16,000 × *g*, 4 °C), and the supernatant was used for protein purification using an ÄKTA Pure system (Cytiva, Sweden). The cleared lysate was loaded onto a 1 mL Ni²⁺-charged HisTrap column (Cytiva, Sweden, #17524701), and proteins were eluted by gradually increasing imidazole concentration in buffer (50 mM Tris-HCl, pH 7.5, 500 mM NaCl, 500 mM imidazole). To remove imidazole, the eluates were desalted on 5 mL desalting Superdex G-25 columns (Cytiva, Sweden, #17140801) into 20 mM Tris-HCl (pH 7.5), 200 mM NaCl.

For TRIP4_ASCH, ZmASCH, and TthASCH, an additional heparin affinity chromatography step was performed. IMAC-purified and desalted proteins were loaded onto a 1 mL HiTrap^TM^ Heparin HP column (Amersham Biosciences, Sweden, #17-0406-01) that was pre-equilibrated with 50 mM Tris-HCl, pH 7.5, 200 mM NaCl. Proteins were eluted by gradually increasing the NaCl concentration in the buffer up to 1 M. The target elution fractions were collected and subjected to an additional desalting step.

Protein purity was analyzed using the standard SDS-PAGE protocol (51), and concentrations were determined using Qubit™ 4 fluorometer (Invitrogen), with the Qubit™ Protein BR Assay Kit (Invitrogen, USA, #A50668) according to the manufacturer’s recommendations.

### Generation and purification of AAAD_HTH fusion proteins

The YqfB-AAAD_HTH fusion proteins were generated by overlapping PCR with 2× Phusion Plus PCR Master Mix (Thermo Scientific, Lithuania, #F631L) to fuse the C-terminal domains of Afu3ASCH and Pku4ASCH (residues 100-177 and 104-190, respectively) to the full-length YqfB protein. In the first step, separate PCR reactions were performed to amplify the two fragments using primers with a 20 nt overlap. In the second step, the purified fragments were used as templates for an overlap extension PCR. After 10 rounds of amplification, primers designed for ligation-independent cloning into a pLATE51 vector (aLICator LIC Cloning and Expression Kit 2, Thermo Fisher Scientific, Lithuania, #K1251) were added to amplify the full-length fusion product. The sfGFP-AAAD_HTH proteins were generated using the same method, however, the protein genes were overlapped by introducing a recognition sequence for the WELQut protease. All DNA primers used for protein fusion are listed in Supplementary Table S3.

To isolate the nucleic acid–binding C-terminal HTH domains of Afu3ASCH and Pku4ASCH, sfGFP–AAAD_HTH fusion proteins were first purified by IMAC, as described above, and subsequently incubated with WELQut protease (Thermo Scientific^TM^, Lithuania, #EO0861) for 12 h at 30°C. Prior to loading onto a 1 mL CM Sepharose Fast Flow column (Cytiva, Sweden, #17071910), the samples were diluted 5-fold with 20 mM Tris-HCl (pH 7.5) buffer to reduce the NaCl concentration for ion-exchange chromatography. The target HTH domains were eluted by gradually increasing NaCl concentration up to 1 M. The target elution fractions were collected and subjected to an additional desalting step.

### Enzymatic activity assay

The enzymatic activity of ASCH domain-containing proteins toward ac^4^C nucleoside was assessed by thin-layer chromatography (TLC). Reaction mixtures contained 4 mM ac^4^C (Combi-Blocks, USA, #QB-9019), 20 mM Tris-HCl (pH 7.5), 200 mM NaCl, and 1 μM of recombinant protein. After incubation at 30°C for 1, 4, or 20 h, 1 μL of each reaction mixture was spotted on silica gel-coated 60 F254 aluminum plates (Merck, USA, #1055540001) and separated using a chloroform:methanol mixture (5:1 v/v). Dried plates were visualized under short-wave UV light (254 nm). Cytidine (Acros Organics, Belgium, #111810500) was used as a positive control.

### Electrophoretic mobility shift assay (EMSA)

Electrophoretic mobility shift assays (EMSA) were performed to assess binding of proteins to RNA and DNA substrates. Unless stated otherwise, binding reactions were carried out at 22°C for 30 min in a buffer containing 20 mM Tris-HCl (pH 7.5), 100–300 mM NaCl, 10% glycerol, and 100 ng/µL bovine serum albumin (BSA).

For EMSA experiments, total *E. coli* MRE 600 tRNA was purchased from Roche (Switzerland, #10109541001). Short RNA and DNA oligonucleotides (ssRNA 17 nt, ssRNA 30 nt, and ssDNA 30 nt) were described previously (43), synthesized by Azenta Life Sciences (Germany), and are listed in Supplementary Table S5. All substrates were dissolved in ultrapure MilliQ water treated with diethylpyrocarbonate (DEPC; Carl Roth, Germany, #K028) and 5′-end labeled with [γ-³²P]-ATP using T4 polynucleotide kinase (Thermo Scientific, Lithuania, #EK0031). Radiolabeled substrates (1 nM) were incubated with 100 nM target proteins (substrate:protein ratio 1:100). Following incubation, samples were resolved on 10% (v/v) native polyacrylamide gels in 0.5× TBE buffer at 200 V for 120 min and visualized using a phosphorimaging system (FLA-5100, Fujifilm, Japan).

Single-stranded 24-nt DNA and its duplex form were used as substrates as described previously (40). The 5′-FAM–labeled ssDNA_24mer was annealed with the complementary strand (ssDNA_24mer_Rv) at a molar ratio of 1:1.5 by heating to 95°C for 5 min followed by gradual cooling to 22°C. EMSA reactions contained 10 nM single- or double-stranded DNA and 500 nM protein (DNA:protein ratio 1:50) and were incubated under the conditions described above. Samples were resolved on 10% (v/v) native polyacrylamide gels in 0.5× TBE buffer at 150 V for 30 min. Fluorescent signals were detected at 435 nm using the FLA-5100 fluorescent image analyzer (Fujifilm, Japan).

### Preparation of bulk tRNA

Bacteria transformed with ASCH domain-containing protein gene-carrying plasmids were inoculated from overnight cultures into 50 mL of sterile LB medium supplemented with either 50 μg/mL ampicillin or 40 μg/mL kanamycin, ensuring a starting OD_600_ of 0.02. Bacteria were grown for about 3 h at 37°C until culture OD_600_ reached ∼0.6. Then, 0.5 mM IPTG and 0.1% rhamnose were added for the KRX strain to initiate protein synthesis. Induction was carried out for 4 h at 30°C. The cells were collected by centrifugation and frozen overnight. Synthesis of soluble recombinant proteins was analyzed by standard SDS-PAGE, and in several cases, by Western blot. Results are presented in Supplementary Figure S2.

The bulk *E. coli* tRNA was extracted using the phenol-chloroform precipitation technique and then purified by ion exchange chromatography with a HiTrap DEAE Sepharose FastFlow column (Cytiva, Sweden, #17505501) and an AKTA Pure FPLC system, as described previously (7).

### Western blot analysis

In short, 1 mL of induced bacterial cultures (see “Preparation of bulk tRNA”) was harvested by centrifugation and resuspended in buffer containing 20 mM Tris-HCl (pH 7.5) and 200 mM NaCl. Cells were lysed using an ultrasonic disintegrator, followed by centrifugation at 30 000 × *g* for 5 min at 4°C to remove cell debris and insoluble protein aggregates. Soluble protein fractions were mixed with denaturing loading dye, heat-denatured, and separated by standard SDS-PAGE on a 14% polyacrylamide gel (Rotiphorese® Gel 40 (29:1), Carl Roth, Germany, #A515.1), using a Bio-Rad Mini-PROTEAN vertical electrophoresis system (Bio-Rad, USA, #1658000EDU). PageRuler™ Plus Prestained Protein Ladder (Thermo Scientific, Lithuania, #26620) was used as a protein molecular size standard.

Following electrophoresis, proteins were transferred onto a 0.45 μm nitrocellulose membrane (Thermo Scientific, USA, #88018). The membrane was blocked for 1 h at 22°C in 2% bovine serum albumin solution prepared in Tris-buffered saline with Tween 20 (TBS-T: 10 mM Tris-HCl, pH 7.5, 150 mM NaCl, 0.1% Tween 20). The membrane was then incubated overnight at 4°C with mouse monoclonal anti-6×His tag primary antibody (Thermo Fisher Scientific, USA, #MA1-21315), diluted 1:1000 in the blocking solution. Afterwards, the membrane was thoroughly washed with TBS-T and incubated for 1h at 22°C with horseradish peroxidase (HRP) conjugated goat anti-mouse secondary antibody diluted 1:10000 (Carl Roth, Germany, #3KY5.1). Protein-antibody complexes were visualized using Pierce^TM^ enhanced chemiluminescence (ECL) blotting substrate (Thermo Scientific^TM^, USA, #32209) and captured using Azure C280 imaging system (Azure Biosystem, USA).

### Enzymatic tRNA hydrolysis

For the enzymatic tRNA hydrolysis, 30 μg (∼10.5 μM) of purified tRNA was heat denaturated for 2 min at 95°C, rapidly cooled down and digested in 50 mM Tris-HCl, pH 8, 10 mM MgCl_2_ with 250 U of Pierce Universal Nuclease (Thermo Scientific^TM^, Lithuania, #88702), 0.005 U Snake Venom Phosphodiesterase from *Crotalus atrox* (Sigma-Aldrich, USA, #P4506), and 2 U of FastAP Thermosensitive Alkaline Phosphatase (Thermo Scientific^TM^, Lithuania, #EF0654) for 4 hours at 37°C. The reactions were terminated by addition of an equal volume of acetonitrile, followed by heating for 10 min at 37°C. Protein precipitates were removed by centrifugation at 30130 × *g* for 10 min at 4°C.

### Evaluation of ac^4^C levels in total tRNA

5 μL of supernatant were analyzed using a Nexera X2 UHPLC liquid chromatography-tandem mass spectrometry (LC-MS/MS) system, coupled with an LCMS-8050 mass spectrometer (Shimadzu, Japan), equipped with an ESI source. Hydrolysis products were separated using a 3 × 150 mm YMC-Triart C18 column (YMC, Japan, #TA12S03-1503WT). Samples were loaded in an aqueous solution containing 0.1% formic acid (solvent A) and eluted using a gradient of increasing acetonitrile concentration (solvent B). The following elution program was used: isocratic 5% B for 1 min, 5% to 95% B over 5 min, isocratic 95% B for 2 min, 95% to 5% B over 1 min, and isocratic 5% B for 4 min. Modified nucleosides were detected using the following ion transitions: *m/z* 247 → 84 for dihydrouridine (D) and 284 → 241 for ac^4^C. Data analysis was performed using LabSolutions LCMS v5.82 SP1 software (Shimadzu, Japan). The relative amounts of ac^4^C in samples were quantified by normalizing the ac^4^C peak areas to the D, enabling sample-to-sample comparison.

### Enzyme activity assay with tRNA in vitro

The activity of purified ASCH domain-containing proteins was tested in vitro with total tRNA from *E. coli* MRE600 (Roche, Switzerland, #10109541001) and tRNA (type V) from wheat germ (Sigma-Aldrich, Germany, #R7876). The 100 μL reaction consisted of 100 μg (∼35 μM) of tRNA, 5 μM of ASCH protein, 20 mM Tris-HCl, pH 7.5, and 200 mM NaCl. After 4 h at 30°C, the reaction was terminated by adding an equal volume of acetonitrile. After centrifugation, the supernatant was mixed with 2.5 volumes of ethanol for tRNA precipitation. The tRNA was enzymatically digested into nucleosides and analyzed by HPLC-MS/MS as described above.

### Phylogenetic analysis

Approximately 16,000 ASCH-domain sequences were obtained from InterPro (52). These sequences were clustered using CD-HIT (53) at a 75% sequence identity threshold, resulting in a nonredundant dataset of 6,282 sequences. Three-dimensional structures for all sequences were predicted using OmegaFold (54). All models were aligned to the *E. coli* YqfB structure with TM-align (55), and only sequences yielding an alignment of at least 80 amino acids according to TM-align were retained for tree construction. From each alignment, a cropped structural model corresponding to the region best matching YqfB was generated.

A structural distance matrix was constructed by pairwise alignment of the cropped structures, extraction of TM-scores, and definition of structural distance as 100 × (1 − TM-score). This distance matrix was then used to build an UPGMA tree using Biopython (56). The resulting tree containing 6,282 leaves is too large for meaningful visualization. Therefore, 500 leaves were randomly selected to generate representative trees (the procedure was repeated several dozen times to ensure preservation of overall topology), and proteins of interest were highlighted for visualization using ETE3 (57).

Structures of the C-terminal domains of Afu3ASCH, Pku2ASCH, and Pku4ASCH, as well as the nucleic acid-binding domains of selected proteins with known nucleic acid-binding functions, were created by using Alphafold3 (58). Then, structural similarity was inferred by using the function ‘All against all’ of the DALI server (59). The similarity was visualized by using the iTOL tool (60).

### Statistical analysis

All experiments were performed with three independent biological replicates. Data are presented as mean ± standard deviation (SD). Differences among multiple groups were assessed by one-way analysis of variance (ANOVA), followed by Dunnett’s multiple comparison test to compare each experimental group with the corresponding control. All statistical analyses were performed using GraphPad Prism 10 for macOS version 10.6.0 (GraphPad Software, USA, www.graphpad.com).

## RESULTS

Currently, the InterPro database (52) contains about 16,000 protein entries that include an ASCH domain. In the initial bioinformatic analysis of the ASCH protein family (41), ASCH domain-containing proteins were classified into 10 subgroups based on sequence features and were shown to share a conserved eight-amino acid motif, G*K**[E/T/S]*R, referred to as the ASCH domain-specific octapeptide (41). Despite extensive amino acid sequence variability across this superfamily, members generally adopt a conserved β-barrel/α-helices fold. To provide an overview of the structural divergence within the ASCH protein family, a structure-based phylogenetic tree was constructed, consisting of 500 randomly selected ASCH family members (the procedure was repeated several dozen times to ensure preservation of overall topology) from all life domains and viruses, to represent the major phylogenetic clades (Figure 1a). ASCH domain-containing proteins clustered into three main subgroups regarding the composition of the domain-specific octapeptide. The subgroup G*K**E*R is commonly found within the ASCH superfamily, and includes members from bacteria (ZmASCH), archaea, eukaryotes (EOLA1, TRIP4), and viruses (45, 46, 49, 50). The less common octapeptide G*K**[T/S]*R, found in YqfB-type ac^4^C amidohydrolases (38, 39), is shared between bacteria, archaea, and some eukaryotes. Non-standard G*K**E*(not R) octapeptide-containing proteins are also common, and such combination is enriched in bacteria. Diversification at the terminal position of the octapeptide, along with its almost exclusive distribution across bacteria, suggests independent evolutionary adaptation events.

**Figure 1:**
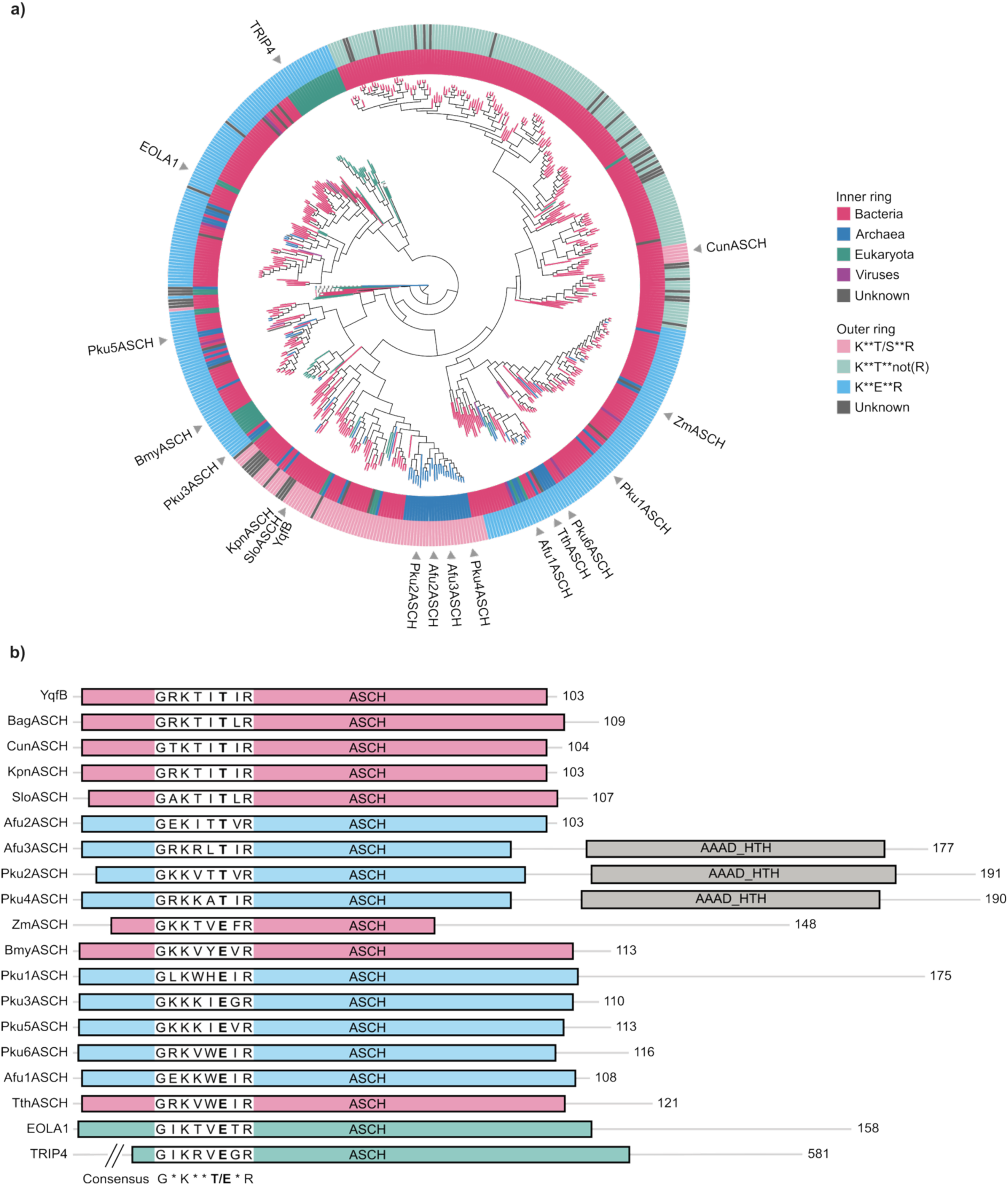
Phylogenetic analysis of ASCH domain-containing proteins. **a)** structure-based phylogenetic tree of 500 ASCH domain-containing representatives. The inner ring represents taxonomic distribution, while the outer ring represents octapeptide variations within the family. **b)** Schematic representation of the target protein organization: *E. coli* (YqfB), *Butiauxella agrestis* (Bag), *Cronobacter universalis* (Cun), *Klebsiella pneumoniae* (Kpn), *Shewanella loihica* (Slo), *Bacillus mycoides* (Bmy), *Thermus thermophilus* (Tth), *Zymomonas mobilis* (Zm), *Archaeoglobus fulgidus* (Afu), *Pyrococcus kukulkanii* (Pku), *Homo sapiens* (TRIP4, EOLA1).

Based on sequence variability and phylogenetic relationships, a total of 19 ASCH-containing proteins were selected for further experimental characterization (marked with triangles in Figure 1a). This set represents all major phylogenetic branches and the semi-conserved octapeptide variants, except for bacterial sequences carrying substitutions at the terminal position, which were not included in this study. Some of the chosen proteins had been partially described in previous reports (EOLA1 (46, 47), TRIP4 (45), ZmASCH (49, 50), YqfB-type amidohydrolases (38–40)), while others are newly identified candidates that belong to mesophilic and thermophilic bacteria such as *B. mycoides* (BmyASCH), *T. thermophilus* (TthASCH), or thermophilic archaea *A. fulgidus* (AfuASCH) and *P. kukulkanii* (PkuASCH) (Figure 1b). Proteins were chosen based on both biological relevance and practical considerations: availability of the source organisms (either already maintained in our laboratory or accessible through commercial collections), and the presence of multiple divergent homologs within the same genome. Together, this representative subset reflects the evolutionary and structural landscape of the ASCH domain and facilitates further biochemical and functional characterization.

### ASCH domain-containing proteins decrease ac^4^C content in bacterial and eukaryotic tRNAs

Based on previous findings, YqfB-type mesophilic proteins containing a threonine in their conserved octapeptide motif act as amidohydrolases with high activity toward ac^4^C and various acylated cytidine derivatives. In contrast, it was recently shown that ZmASCH, TRIP4_ASCH, and EOLA1 proteins, which contain a glutamate residue in the same motif, do not hydrolyze ac^4^C (40). To investigate whether this variation within the octapeptide influences nucleoside hydrolysis activity, we tested recombinant proteins against ac^4^C in vitro. Protein activities were analyzed by TLC after 1, 4, and 24 hours of incubation (Figure 2a and Supplementary Figure S1), with YqfB-type amidohydrolases serving as a positive control. The results indicated that hydrolytic activity did not strictly correlate with the octapeptide sequence, since proteins containing either threonine or glutamate could hydrolyze ac^4^C. After 24 hours, ZmASCH and TRIP4_ASCH remained inactive toward ac^4^C, whereas EOLA1 exhibited partial hydrolysis during prolonged incubation. All other tested proteins displayed varying levels of activity: Bmy, Tth, Afu2, Pku1, Pku3, and Pku4 completely hydrolyzed the substrate within the first four hours, while Afu1, Afu3, and Pku5 showed incomplete hydrolysis under the same conditions (Figure 2a).

**Figure 2:**
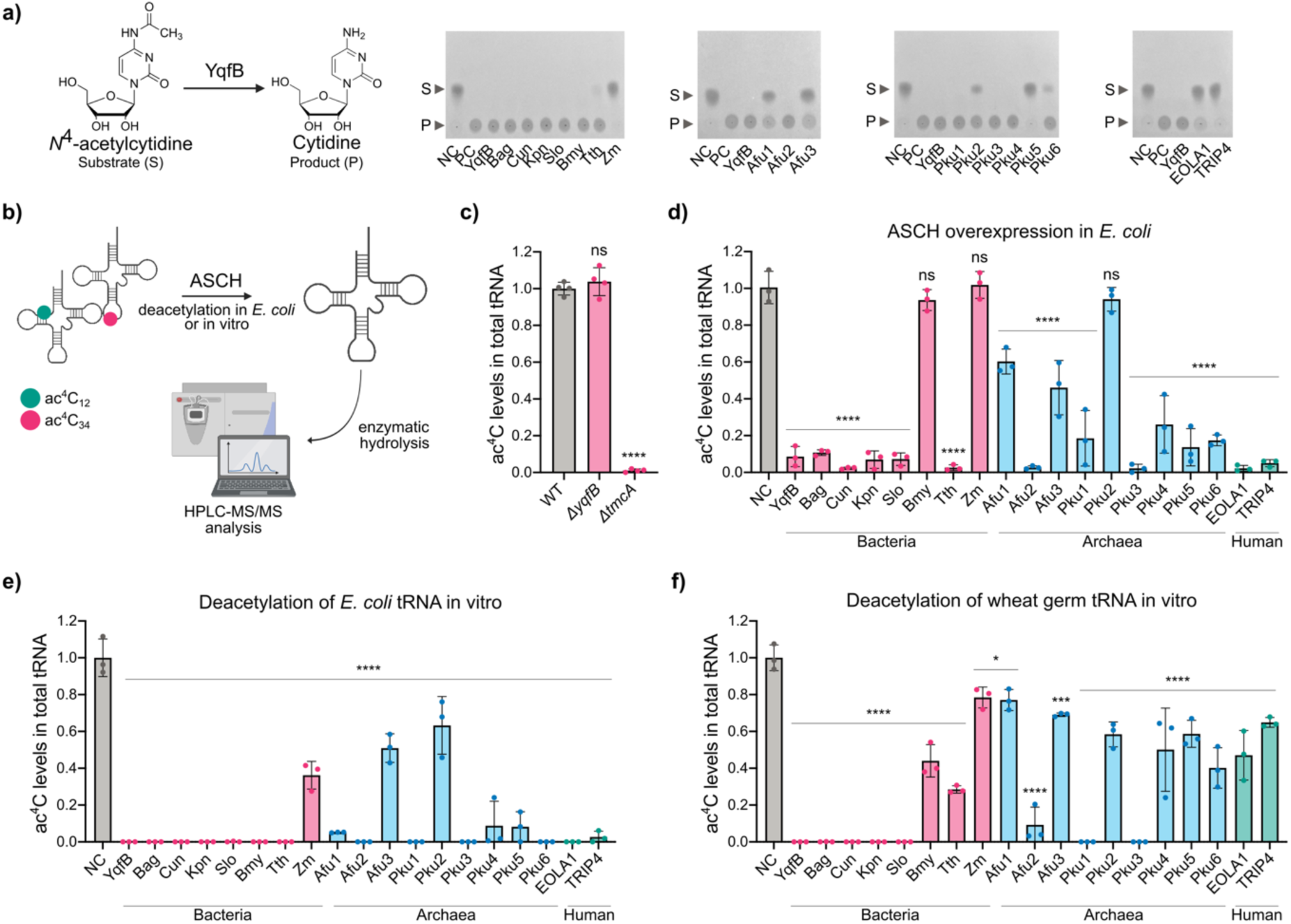
Functional characterization of ASCH domain–containing proteins. **a)** TLC analysis of the amidohydrolytic activity of ASCH domain-containing proteins toward ac^4^C after 4 hours of incubation. S – substrate (ac^4^C), P – product (cytidine), NC – negative control/substrate, PC – positive control/product. **b)** Experimental workflow of ac^4^C level determination in bacterial and eukaryotic total tRNA by HPLC-MS/MS. **c)** HPLC-MS/MS analysis of endogenous YqfB activity in *E. coli* BW25113 strain and its single-gene knockout mutants; **d)** ac^4^C levels in total tRNA of *E. coli* KRX strain upon the overexpression of ASCH domain-containing proteins. Activity of recombinant ASCH domain-containing proteins in vitro using **e)** *E. coli* MRE600 total tRNA or **f)** wheat germ total tRNA. **** *p* < 0.0001, *** *p* < 0.001, * < 0.05, ns *p* ≥ 0.05 compared to negative control (empty vector), one-way ANOVA with Dunnett’s post hoc test.

Since ac^4^C in bacteria is predominantly associated with tRNA, the target ASCH domain-containing proteins were tested for tRNA deacetylase activity both upon induction in *E. coli* and in vitro (Figure 2b). To begin with, we examined whether the endogenous *E. coli* YqfB amidohydrolase activity could be detected at the tRNA level. For this, the tRNA fraction purified from wild-type *E. coli* BW25113 strain cells served as a reference control in HPLC-MS/MS analysis, while tRNA fractions from KEIO collection single-gene knockout strains BW25113 *ΔyqfB* and BW25113 *ΔtmcA* were analyzed in parallel. The results, shown as a normalized ratio of total ac^4^C and dihydrouridine (D, internal control) levels, revealed no significant differences between the wild-type BW25113 and its *ΔyqfB* strain, whereas the deletion of the modification writer TmcA left only trace amounts of ac^4^C (Figure 2c). These results support an earlier report, which also showed no significant differences in ac^4^C levels upon YqfB deletion (40). We suggest that under standard growth conditions, cells maintain a stable ac^4^C abundance through constitutive TmcA activity, whereas the activity of potential erasers, such as YqfB, could be condition-dependent and tightly regulated.

Although no significant changes were observed in the native *E. coli* strains, we further evaluated the possible ac^4^C tRNA eraser activity of the ASCH domain by overexpressing each target protein in *E. coli* KRX cells for 4h. The empty expression vector control (NC) represents total modification levels, presented as normalized ac^4^C/D ratio in purified tRNA fraction, in the presence of endogenous TmcA and YqfB. Most proteins significantly reduced ac^4^C levels, and some even maintained them comparable to those of the *ΔyqfB* strain (YqfB-type amidohydrolases, TthASCH, Afu2ASCH, Pku3ASCH, EOLA1, TRIP4_ASCH), indicating effective removal of the modification despite constitutive TmcA activity. The results revealed that only BmyASCH, ZmASCH, and Pku2ASCH showed no significant decrease in the ac^4^C levels (Figure 2d).

Furthermore, to eliminate the reversible effect of the native modification writer TmcA, the purified proteins were tested in vitro with total tRNA from *E. coli* MRE600 and wheat germ (Figure 2e, f). This analysis revealed that all tested ASCH domain-containing proteins removed the bacterial ac^4^C34 modification, most leaving only trace amounts of detectable nucleoside, indicating direct enzymatic tRNA deacetylation (Figure 2e). Even proteins that failed to alter ac^4^C levels upon expression in *E. coli* were active in vitro, suggesting that their activity in the cell is masked by the activity of TmcA. In contrast, the efficiency of enzymatic demodification of eukaryotic tRNA was more variable: YqfB-type mesophilic amidohydrolases, Afu2ASCH, Pku1ASCH, and Pku3ASCH, although sequence- and structure-wise divergent, readily removed the ac^4^C12 modification from the double-stranded region of the D-stem in vitro. Other proteins, although still capable of removing the acetyl group, were less effective, suggesting divergent substrate recognition patterns within the protein family (Figure 2f).

### Conserved residues within the ASCH octapeptide motif are essential for catalysis

ASCH domain-containing proteins can vary greatly in their amino acid sequence yet maintain a highly conserved β-barrel fold. We selected the 121 amino acid ASCH protein from *T. thermophilus* HB8 (TthASCH; PDB 2DP9) as a representative of the glutamate-containing ASCH octapeptide subgroup (G*K**E*R), in contrast to the threonine-containing YqfB-type variants (G*K**T*R). The amino acid sequence overlap between YqfB and TthASCH is low, at only 27%; however, both proteins adopt a characteristic β-barrel conformation. Therefore, to investigate the alternative catalytic mechanism of TthASCH, five candidate amino acid residues were selected based on the previously suggested YqfB catalysis mechanism and 3D structural superimposition of TthASCH (2DP9) and ac^4^C-containing chain B of YqfB (9KYF) crystal structure model (Figure 3a). By employing a site-directed mutagenesis approach, the following mutant variants were obtained: Tyr14Ala, Tyr14Phe, Lys23Ala, Lys23Leu, Glu26Ala, Glu26Asp, Glu26Ser, Glu26Thr, Arg28Ala, Arg28Leu, Tyr81Ala, Tyr81Phe, and Tyr81Leu. All mutant variants were purified, and their activity towards ac^4^C was analyzed by applying TLC (Figure 3b). The amidohydrolytic activity of TthASCH was disrupted in all variants except Tyr14Leu, Glu26Asp, Tyr81Phe, and Tyr81Leu, which retained partial activity. This indicates that an acidic residue at position Glu26 and hydrophobic contacts at Tyr14 and Tyr81 are necessary for nucleoside positioning and hydrolysis. Additionally, since the reconstruction of a YqfB active site by introducing threonine instead of glutamate was not enough to maintain hydrolytic activity, this suggests an alternative nucleoside positioning and hydrolysis mechanism in TthASCH. The effects of these point substitutions on TthASCH activity were further tested by overexpressing mutant variants in the *E. coli* KRX strain. In contrast to the results obtained using free nucleoside, the activity of TthASCH mutants on tRNA revealed a catalytically essential dyad comprising Lys23 and Glu26 (Figure 3c).

**Figure 3:**
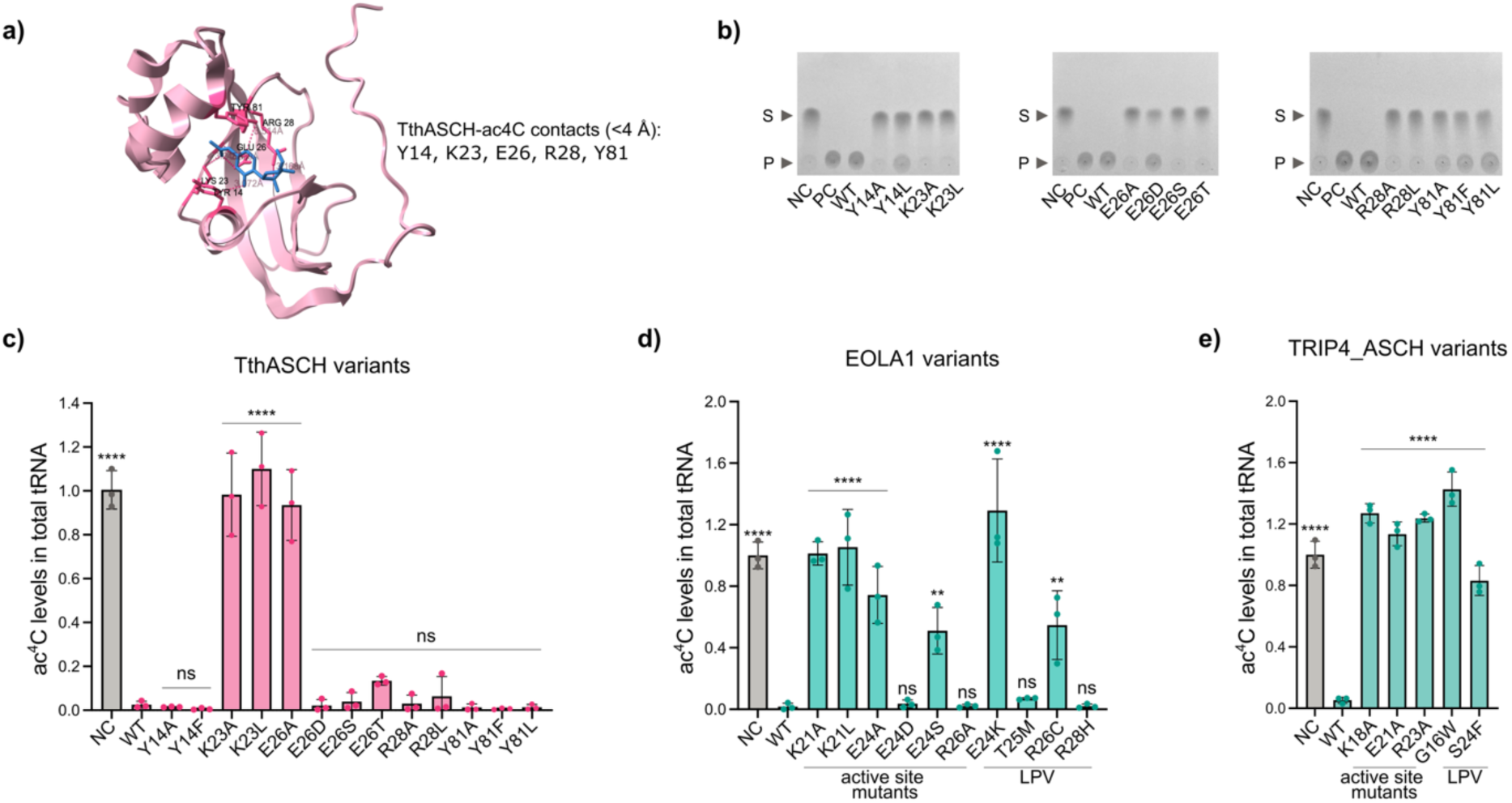
Characterization of G*K**E*R octapeptide-containing proteins. **a)** YqfB-ac^4^C crystal structure (9KYF) based identification of catalytic residues of TthASCH (2DP9). **b)** TLC analysis of the effect of single point mutations of TthASCH on the amidohydrolytic activity towards ac^4^C. ac^4^C levels in total tRNA of *E. coli* KRX strain upon the overexpression of **c)** TthASCH mutant variants, **d)** EOLA1 mutant variants, and **e)** TRIP4_ASCH mutant variants. **** *p* < 0.0001, *** *p* < 0.001, ** *p* < 0.01, * *p* < 0.05, ns *p* ≥ 0.05 compared to wild-type (WT) protein, one-way ANOVA with Dunnett’s post hoc test. LPV – likely pathogenic variants.

We next assessed potential catalytic amino acids in the human ASCH domain-containing proteins EOLA1 and TRIP4_ASCH, also belonging to the octapeptide subgroup G*K**E*R. To evaluate how the ASCH octapeptide residues (lysine, glutamate, and arginine) affect tRNA demodification, we generated point mutants, overexpressed them in *E. coli*, and compared the resulting ac^4^C levels with those of the corresponding wild-type proteins. The results revealed that in both EOLA1 (Figure 3d) and TRIP4_ASCH (Figure 3e), substitutions of the conserved lysine residue led to catalytically inactive variants, resembling the essential role of lysine in YqfB-type amidohydrolases and TthASCH. Glutamate-to-alanine substitutions were also found to be critical for protein activity. However, the glutamate-to-aspartate substitution was well tolerated in EOLA1, resulting in ac^4^C levels comparable to the wild-type enzyme. The differences between the two proteins became apparent when arginine was replaced with alanine: the EOLA1 mutant showed no significant loss of activity, whereas the analogous TRIP4_ASCH mutant was catalytically inactive. These results indicate that lysine and glutamate/threonine are crucial residues for tRNA deacetylation, while the importance of arginine might be protein specific.

We also investigated how naturally occurring point mutations may impact the catalytic function of these enzymes. According to the UniProt (61) Variant Viewer, multiple non-synonymous substitutions in EOLA1 and TRIP4_ASCH have been predicted to generate likely pathogenic variants (LPVs). To test whether the loss of tRNA deacetylase activity could lead to the predicted pathogenic outcomes, we selected several predicted positions that are closely located near the catalytic residues. In EOLA1, the Thr25Met and Arg28His variants showed no detectable changes in ac^4^C levels compared to the wild-type enzymes, indicating minimal impact on catalytic function (Figure 3d). By contrast, Arg26 substitution with cysteine resulted in a ∼70-fold increase in relative ac^4^C levels, and the Glu24Lys mutant displayed a complete loss of detectable ac^4^C hydrolysis activity. LPV-associated TRIP4_ASCH variants, including Gly16Trp and Ser24Phe, both resulted in a complete loss of detectable deacetylase activity compared with the wild-type protein (Figure 3e). Together, these findings suggest that several disease-linked mutations in human ASCH domain-containing proteins affect their ability to remove ac^4^C from tRNA, potentially linking disrupted RNA deacetylation to pathogenic phenotypes.

### Determination of nucleic acid binding activity

In our previous study, no nucleic acid–binding activity was detected for mesophilic YqfB-type amidohydrolases (38), consistent with the recently updated crystal structure of *E. coli* YqfB (40). In contrast, several target ASCH domain-containing proteins analyzed in this work possess a net positive charge, suggesting potential nucleic acid-binding capabilities. Notably, TthASCH, Afu1ASCH, Afu2ASCH, Afu3ASCH, Pku2ASCH, Pku4ASCH, Pku5ASCH, and Pku6ASCH exhibit isoelectric points comparable to ZmASCH and TRIP4-ASCH (close or above pI 9), both of which were previously shown to interact with nucleic acids (45, 49).

Given that ac⁴C modification is predominantly associated with specific tRNA species in prokaryotes, we first examined the ability of these proteins to bind tRNA using γ-³²P–labeled *E. coli* MRE600 total tRNA in electrophoretic mobility shift assays (EMSA). Under standard conditions containing 100 mM NaCl, TthASCH, Afu3ASCH, Pku2ASCH, Pku4ASCH, and TRIP4-ASCH exhibited clear tRNA-binding activity that was independent of divalent metal ions (Figure 4a). Increasing the ionic strength up to 300 mM NaCl reduced binding, and only TthASCH, Afu3ASCH, and Pku4ASCH retained detectable tRNA association (Figure 4b). Interestingly, Afu3ASCH and Pku4ASCH produced additional slower-migrating complexes, suggesting possible multimerization of these proteins on single tRNA molecules or binding to specific structural motifs within the molecule.

**Figure 4:**
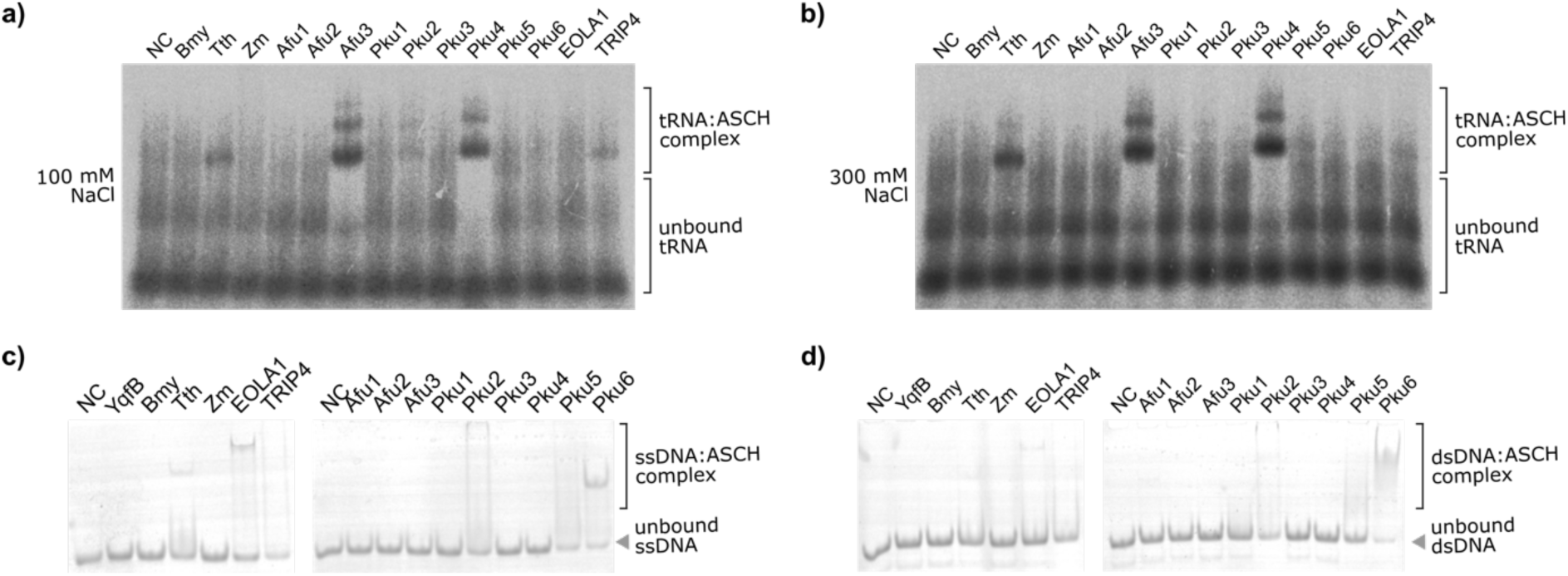
EMSA analysis of ASCH domain-containing proteins nucleic acid binding abilities: **a)** *E. coli* MRE600 total tRNA binding by target ASCH proteins in 100 mM NaCl; **b)** *E. coli* MRE600 total tRNA binding by target ASCH proteins in 300 mM NaCl. **c)** binding to 24 nt ssDNA primer; **d)** binding to 24 nt DNA duplex.

To further assess substrate specificity, we evaluated protein binding to short nucleic acid oligonucleotides using 5’-FAM-labeled 24 nt ssDNA/dsDNA substrates as listed in (37). EMSA results demonstrated the formation of protein:DNA complexes for TthASCH, EOLA1, Pku2ASCH, Pku5ASCH, and Pku6ASCH, indicating that ASCH family proteins exhibit divergent preferences for nucleic acid types or structural motifs. This hypothesis is further strengthened in the case of TthASCH, which exhibits a clear shift of tRNA and ssDNA, whereas only a weak shift of dsDNA was detected. In contrast, no tRNA shifts were detected in the case of Pku6ASCH, however, the protein was found to bind both single and double-stranded DNA. Notably, despite its overall negative charge at pH 7.5 (pI 6.4), EOLA1 was able to bind DNA oligonucleotides but not tRNA, likely owing to the presence of a positively charged nucleic acid–binding region within its structure.

### Structural analysis reveals a novel HTH-type nucleic acid binding domain

To further investigate structural features responsible for nucleic acid interactions, we performed comparative analyses of protein architectures. These proteins fell into two major groups: (i) a TRIP4/ZmASCH-like group comprising proteins TthASCH, Pku5ASCH, Pku6ASCH, and (ii) a group characterized by an additional C-terminal domain, including proteins Afu3ASCH, Pku2ASCH, and Pku4ASCH. The members of the first group have a positively charged cleft associated with nucleic acid binding, however, the second group is of particular interest, as the C-terminal domain does not correspond to any annotated domain in InterPro/Pfam (52) and SMART protein databases (62). A sequence-based homology search (HHpred) (63) revealed the domain’s similarity to bacterial RNA polymerase sigma factors E and H, suggesting a novel structural motif. Further structural analysis of 3D protein models of Afu3ASCH, Pku2ASCH, and Pku4ASCH using the DALI server (59) identified a nucleic acid binding helix-turn-helix (HTH) like conformation, commonly found in the N-terminal domain of LysR-type transcription regulators (e.g. CysB (PDB 9f14), OxyR (PDB 6g1b)). Although the results showed low sequence identity (less than 20%), the structural match is considered strong enough to suggest a potential evolutionary or structural relationship.

To clarify the structural relationships and gain insight into a possible role of the C-terminal domain of archaeal ASCH proteins, we performed a structural comparison between these proteins and selected proteins with a known capability of DNA or RNA binding (Figure 5). Representative proteins were selected having DNA-binding domains (helix-turn-helix motif and winged helix-turn-helix motifs) and RNA-binding domains (PUA, Ferredoxin-fold anticodon binding domain and RNA recognition motif). Additionally, nuclear acid-binding proteins with significant sequence similarity to the C-terminal domains of ASCH (Ring-type E3 ubiquitin transferase, RNA polymerase factors sigma E and H) were included in structural comparison, which was performed by using the function ‘All against all’ of DALI server. Apparently, the C-terminal domains of archaeal ASCH proteins clustered into a single distinctive group (Figure 5) and were structurally similar to LysR_HTH (IPR000847) and winged HTH-like DNA-binding (IPR036388) domain-containing proteins. Despite being structurally related to DNA-binding HTH-like domains, the C-terminal domain of archaeal ASCH proteins might represent a previously unrecognized structural family. We therefore designate this domain as the archaeal ASCH domain–associated HTH (AAAD_HTH) domain.

**Figure 5.**
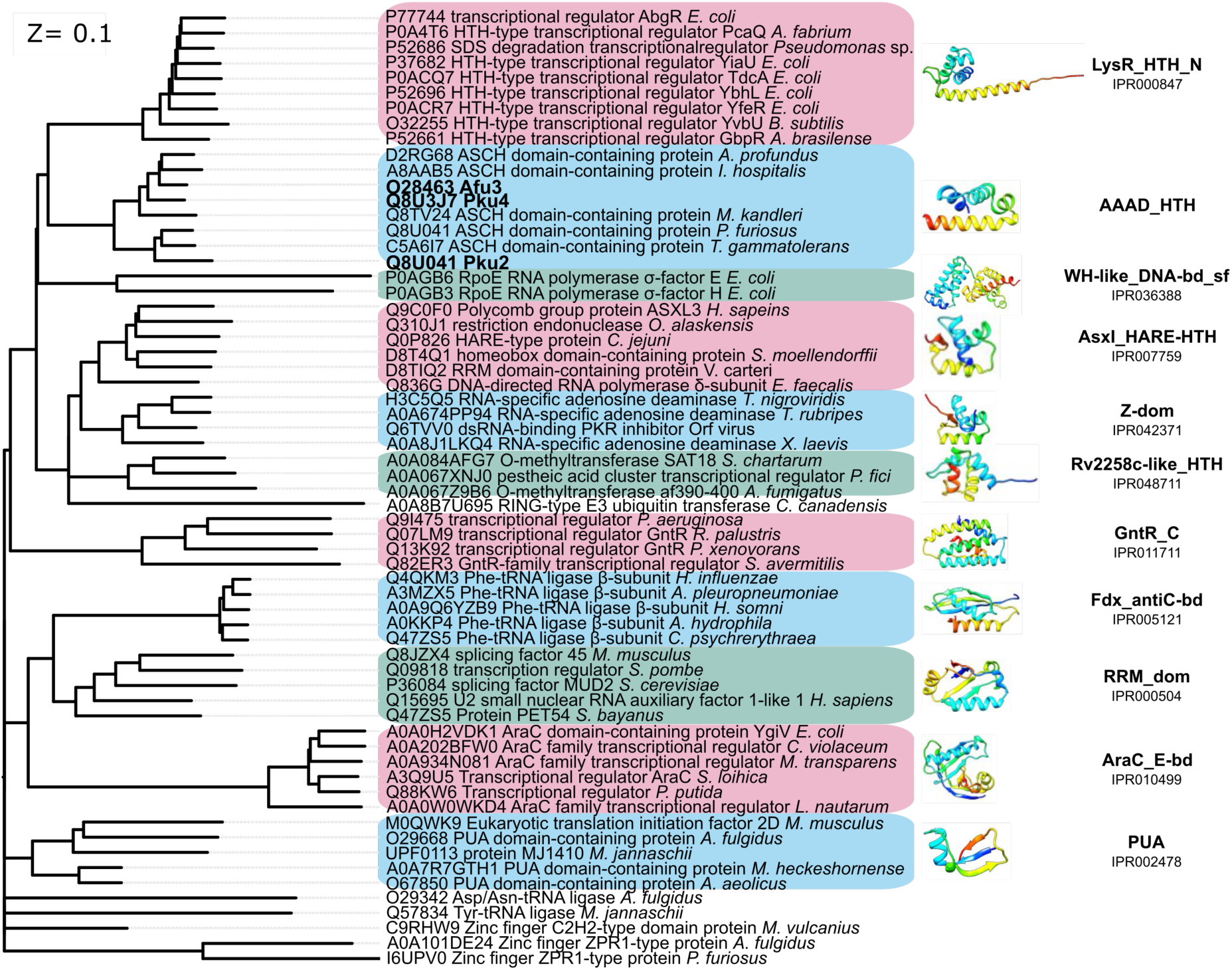
Structural similarity dendrogram of different nucleic acid-binding proteins and C-terminal HTH domains of archaeal ASCH proteins. The dendrogram is derived by average linkage clustering of the structural similarity matrix (Dali Z-scores). Selected proteins are named by Uniprot ID. Different colors represent separate clusters of nucleic acid-binding domains. Domain names and codes were retrieved from the InterPro database.

To test if the AAAD-HTH domain is responsible for interactions with nucleic acids (Figure 6a), the C-terminal HTH domains of Afu3ASCH and Pku4ASCH were joined to nucleic acid non-binding sfGFP and *E. coli* YqfB (Figure 6b). EMSA analysis revealed clear tRNA band shifts by the full length Afu3ASCH protein, the stand-alone AAAD_HTH domain of Afu3ASCH, and the YqfB-Afu3HTH fusion protein, while tRNA binding activity was not detected for the stand-alone catalytic ASCH domain of Afu3ASCH (Figure 6c). Comparable pattern was obtained for the homologous Pku4ASCH constructs, as shown in Supplementary Figure 3a. These results indicate that the AAAD_HTH domain is both necessary and sufficient for concentration-dependent tRNA binding (Supplementary Figure 3b).

**Figure 6.**
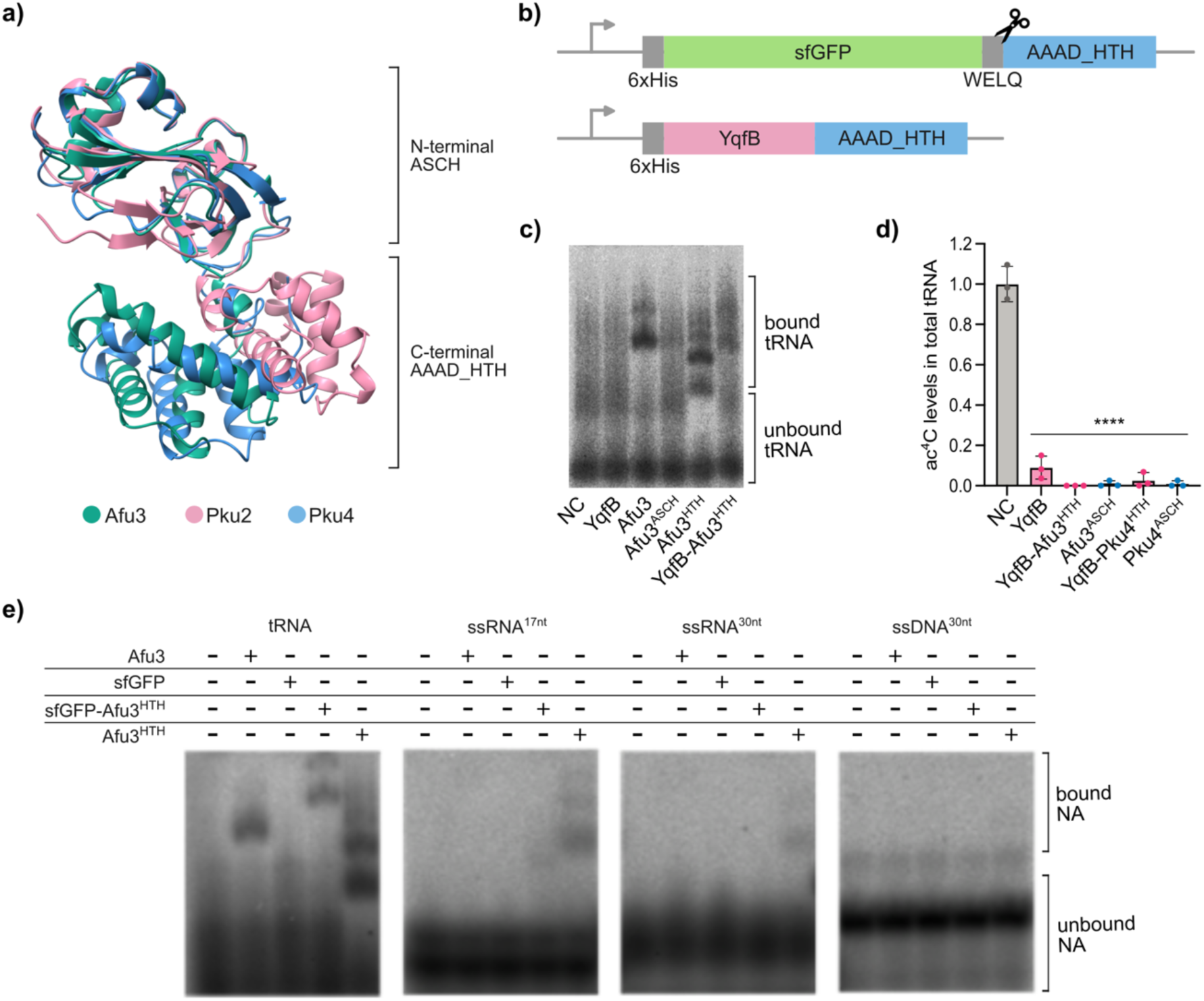
Archaea ASCH domain-associated helix-turn-helix (AAAD_HTH) domain is involved in nucleic acid recognition. **a)** Structural comparison of Alphafold3 models of ASCH proteins containing the AAAD_HTH domain. **b)** Schematic representation of sfGFP-and YqfB-AAAD_HTH fusion protein constructs used in this study. **c)** tRNA binding analysis of Afu3ASCH-derived variants: Afu3 (full-length protein containing both ASCH and HTH domains), Afu3^ASCH^ (stand-alone ASCH domain), Afu3^HTH^ (standalone HTH domain), and YqfB-Afu3^HTH^ fusion protein. **d)** HPLC-MS/MS analysis of ac^4^C levels in total tRNA of *E. coli* KRX strain upon overexpression of a stand-alone ASCH domain of Afu3ASCH and Pku4ASCH, and YqfB-AAAD_HTH fusion proteins. **** *p* < 0.0001, compared to negative control (empty vector), one-way ANOVA with Dunnett’s post hoc test. **e)** Comparison of nucleic acid binding activity of full-length Afu3ASCH and AAAD_HTH-containing protein constructs, including sfGFP (negative control), the sfGFP-Afu3HTH fusion protein, and the stand-alone Afu3^HTH^ domain, using tRNA and ssRNA/ssDNA oligonucleotide substrates.

To assess whether the presence or absence of the AAAD_HTH domain affects enzymatic activity, ac⁴C levels in total tRNA were measured following expression of Afu3ASCH and Pku4ASCH variants lacking the HTH domain in *E. coli* (Figure 6d). Removal of the AAAD_HTH domain from Afu3ASCH or Pku4ASCH did not impair the catalytic activity of the stand-alone ASCH domains. Likewise, fusion of the AAAD_HTH domain to the C-terminus of YqfB did not disrupt its deacetylase activity, demonstrating that the AAAD_HTH domain mediates nucleic acid binding without interfering with catalysis.

## DISCUSSION

In this study, we characterize a set of sequence and structure-divergent ASCH domain-containing proteins from bacteria, archaea, and humans, and evaluate their ability to hydrolyze ac^4^C in both nucleoside and tRNA contexts. The YqfB-type amidohydrolases, which harbor a threonine in their semi-conserved ASCH octapeptide motif, were previously shown to hydrolyze the ac^4^C nucleoside, which was proposed as their primary substrate based on kinetic analyses (38, 39). Recently, several previously characterized ASCH homologs (ZmASCH, EOLA1, TRIP4) carrying a threonine-to-glutamate substitution within the semi-conserved ASCH octapeptide motif were shown to lack ac^4^C deacetylase activity, leading to the proposal that this residue change determines the catalytic function (40). Here we have observed that members of both G*K**T*R (YqfB-type amidohydrolases, Afu2ASCH, Afu3ASCH, Pku2ASCH, Pku4ASCH) and G*K**E*R (BmyASCH, TthASCH, Afu1ASCH, Pku1ASCH, Pku3ASCH, Pku5ASCH, Pku6ASCH, EOLA1) octapeptide motif groups exhibit detectable activity toward ac^4^C both within tRNA and as a free nucleoside. Our findings suggest that more complex sequence or structural features account for the variations in hydrolytic activity, rather than the single change in the semi-conservative octapeptide. The structural and functional divergence within ASCH protein family can be illustrated by the example of YqfB-type ac^4^C amidohydrolases and nucleoside hydrolysis activity-lacking ssRNA nuclease ZmASCH (39, 49). It is not uncommon in nature to reuse existing protein folds and, through evolutionary changes, adapt to divergent functions or different substrate or cofactor specificities (64). For example, it was recently shown that the Rossmann fold containing enzymes can undergo sequence rearrangements through deletions or insertions that preserve domain-specific spatial fold, however, such adaptations change the nucleoside cofactor preferences from NAD to SAM (65).

The nucleic acid-binding diversity we observed within the ASCH family further reflects the evolutionary plasticity of this domain. Although all examined homologs preserve the conserved β-barrel fold, several proteins possess a strongly positive surface charge (pI > 9), which is provided by lysine- and arginine-rich motifs. Such characteristic, typically associated with nucleic acid-binding proteins, can result in less specific charge interactions with the substrate (66, 67). Among the characterized proteins, several interacted with both RNA and DNA molecules, as was previously demonstrated for ZmASCH and TRIP4_ASCH, however, others displayed detectable affinity for only one type of nucleic acid, while some lacked detectable nucleic acid-binding activity despite their overall positive charge (45, 49). Interestingly, despite having an overall negative charge, a previous analysis of the EOLA1 crystal structure suggested possible affinity for RNA, as seen in PUA or YTH domain-containing proteins (68, 69). EOLA1 contains a positively charged cleft that is potentially involved in substrate recognition (40, 46). However, our data show that the protein does not bind tRNA but does interact with single- and double-stranded DNA. In contrast, recent evidence shows that EOLA1 associates with mitochondrial 12S rRNA (48). These observations suggest that the protein may have evolved to recognize multiple nucleic-acid targets or may exhibit sequence- or motif-specific binding. Taken together, our findings suggest that despite sharing a conserved fold, ASCH domain–containing proteins exhibit diversity in nucleic acid recognition. Some homologs show broad affinity for both RNA and DNA, whereas others interact with only one substrate type, or fail to bind any of the tested molecules. Such behavior does not simply align with overall protein charge, as several non-binding proteins are positively charged, while others with a net negative charge contain basic clefts for nucleic acid binding. This diversity suggests functional divergence within the ASCH family, though it remains possible that these proteins could act on cellular substrates not provided by our assay conditions. Notably, an *N*^4^-acetylated 2’-deoxycytidine has recently been identified in the genomic DNA of *Arabidopsis thaliana*, predominantly near transcription start sites (70). These observations raise the possibility that ASCH domain-containing proteins could also have functions in transcriptional regulation.

In a subset of archaeal ASCH homologs, nucleic acid recognition is further regulated by C-terminal HTH domains. HTH domains are usually associated with DNA binding, however, some publications show that these architectures can adapt to bind RNAs (71, 72). Structural phylogeny suggests that ASCH-associated archaeal HTH domains (AAAD_HTH) resemble DNA-binding HTH domains found in bacterial LysR-type transcriptional regulators. These archaeal fusion domains provide a positively charged nucleic acid binding motif, which in Afu3ASCH and Pku4ASCH shows a strong affinity for tRNA and no detectable interactions with the tested DNA substrates. However, although structurally close, Pku2ASCH exhibited weak binding to both tRNA and DNA. The restricted distribution of the archaeal ASCH-associated HTH domain and its distinct binding behavior suggest that it may have adapted to specialized cellular targets.

Despite the differences in structure, nucleoside hydrolysis activity, and nucleic acid recognition, all examined ASCH domain-containing proteins were unified by their ability to remove the ac^4^C modification from tRNAs. Earlier work by Meng et al. reported that *E. coli ΔyqfB* strains display no significant changes in ac^4^C levels in small RNA (<200 nt) and total RNA fractions, and that the elimination of endogenous YqfB does not affect bacterial growth at both standard and elevated temperatures (40). Our current findings support that no detectable ac^4^C differences were found in the tRNA fraction of the BW25113 *ΔyqfB* strain, however this could indicate a specific condition-induced activity within the bacterial cells. Similar observations were also made in the case of 4-thiouridine tRNA erasers – under standard conditions, no significant effect of endogenous RudS desulfidase was detected. However, the gene encoding this protein was found to be located in a light-inducible operon, supporting its function in UV damage control (7). Similarly, only minor growth differences were detected in the knockout strains of bacterial *N*^1^-methyladenosine demethylases RMD1 and RMD2, suggesting that standard laboratory conditions may not provide the optimal environment for these proteins to function (8).

The previously published evidence suggests that higher ac^4^C levels are typically associated with stress conditions, such as heat shock (33). Several research groups have reported increased ac^4^C levels and additional modified positions in the coding and non-coding RNAs of thermophilic archaea (34, 35). Since some archaea possess several genes encoding ASCH domain-containing proteins (*A. fulgidus*, *P. kukulkanii*, and others) that are both structurally and functionally divergent, this suggests that these proteins may have different targets within the cell and could be involved in the dynamic regulation of ac^4^C levels across different RNA species or positions.

However, it should not be overlooked that the function of some ASCH domain-containing proteins may not be directly linked to ac^4^C de-modification, and their catalytic activity could instead be an evolutionary remnant. For example, recent studies indicate that EOLA1 localizes exclusively in mitochondria (48), where tRNAs lack ac^4^C modification (73). It therefore remains unclear whether the deacetylase activity of EOLA1 is physiologically relevant in this context. Notably, in our study, a full-length form of EOLA1, harboring a putative N-terminal mitochondrial localization sequence, was enzymatically active. Together, these observations raise the possibility that EOLA1 either retains a functional deacetylase role in certain contexts or fulfills multiple cellular functions.

To conclude, this study of ASCH domain-containing proteins strengthens the evidence for dynamic regulation of ac^4^C modification and additional regulatory pathways within the cell. While the precise cellular functions of ASCH proteins remain to be determined, the observed differences in biochemical activities point to distinct substrate preferences within this family. In this context, the previous studies demonstrating that the small amidohydrolases YqfB and D8_RL can activate 5-fluorocytidine-derived prodrugs (74) suggest that the diversity of ASCH proteins may be suited to prodrug activation systems with varying specificities. Accordingly, further investigation of the cellular roles and substrate preferences of ASCH family proteins is essential for both fundamental understanding and practical applications.

## ACKNOWLEDGEMENTS

We are grateful to dr. Mindaugas Zaremba for providing *A. fulgidus* genomic DNA. We also thank dr. Justas Vaitekūnas and Gytis Druteika for technical assistance.

Graphical elements from BioRender.com were used to create the graphical abstract and Figure 2b. AI-assisted tools, including Grammarly and ChatGPT, were used solely for language corrections.

## AUTHOR CONTRIBUTIONS

Roberta Statkevičiūtė: Conceptualization, Formal analysis, Investigation, Methodology, Validation, Writing—original draft, Writing—review & editing, Visualization. Mikas Sadauskas: Formal analysis, Methodology, Visualization, Writing—original draft, Writing—review & editing. Agota Aučynaitė: Formal analysis, Funding acquisition, Methodology, Writing—review & editing. Audrius Laurynėnas: Formal analysis, Methodology, Visualization, Writing—original draft, Writing—review & editing. Greta Gakaitė: Investigation, Writing—review & editing. Rolandas Meškys: Conceptualization, Funding acquisition, Resources, Writing—review & editing.

## SUPPLEMENTARY DATA

Supplementary Data are available at NAR online.

## DATA AVAILABILITY

The data underlying this article are available in the article and in its online supplementary material.

## REFERENCES

1. Cappannini, A., Ray, A., Purta, E., Mukherjee, S., Boccaletto, P., Moafinejad, S.N., Lechner, A., Barchet, C., Klaholz, B.P., Stefaniak, F., et al. (2024) MODOMICS: a database of RNA modifications and related information. 2023 update. Nucleic Acids Research, 52, D239–D244.

2. McCown, P.J., Ruszkowska, A., Kunkler, C.N., Breger, K., Hulewicz, J.P., Wang, M.C., Springer, N.A. and Brown, J.A. (2020) Naturally occurring modified ribonucleosides. Wiley Interdiscip Rev RNA, 11, e1595.

3. Liu, F., Clark, W., Luo, G., Wang, X., Fu, Y., Wei, J., Wang, X., Hao, Z., Dai, Q., Zheng, G., et al. (2016) ALKBH1-Mediated tRNA Demethylation Regulates Translation. Cell, 167, 816–828.e16.

4. Li, X., Xiong, X., Wang, K., Wang, L., Shu, X., Ma, S. and Yi, C. (2016) Transcriptome-wide mapping reveals reversible and dynamic N1-methyladenosine methylome. Nat Chem Biol, 12, 311–316.

5. Zheng, G., Dahl, J.A., Niu, Y., Fedorcsak, P., Huang, C.-M., Li, C.J., Vågbø, C.B., Shi, Y., Wang, W.-L., Song, S.-H., et al. (2013) ALKBH5 is a mammalian RNA demethylase that impacts RNA metabolism and mouse fertility. Mol Cell, 49, 18–29.

6. Jia, G., Fu, Y., Zhao, X., Dai, Q., Zheng, G., Yang, Y., Yi, C., Lindahl, T., Pan, T., Yang, Y.-G., et al. (2011) N6-Methyladenosine in nuclear RNA is a major substrate of the obesity-associated FTO. Nat Chem Biol, 7, 885–887.

7. Jamontas, R., Laurynėnas, A., Povilaitytė, D., Meškys, R. and Aučynaitė, A. (2024) RudS: bacterial desulfidase responsible for tRNA 4-thiouridine de-modification. Nucleic Acids Res, 52, 10543–10562.

8. Foo, M., Frietze, L.R., Enghiad, B., Yuan, Y., Katanski, C.D., Zhao, H. and Pan, T. (2024) Prokaryotic RNA N1-Methyladenosine Erasers Maintain tRNA m1A Modification Levels in Streptomyces venezuelae. ACS Chem. Biol., 19, 1616–1625.

9. Thomas, G., Gordon, J. and Rogg, H. (1978) N4-Acetylcytidine. A previously unidentified labile component of the small subunit of eukaryotic ribosomes. Journal of Biological Chemistry, 253, 1101–1105.

10. Zachau, H.G., Dütting, D. and Feldmann, H. (1966) The structures of two serine transfer ribonucleic acids. Hoppe Seylers Z Physiol Chem, 347, 212–235.

11. Kowalski, S., Yamane, T. and Fresco, J.R. (1971) Nucleotide Sequence of the ‘Denaturable’ Leucine Transfer RNA from Yeast. Science, 172, 385–387.

12. Staehelin, M., Rogg, H., Baguley, B.C., Ginsberg, T. and Wehrli, W. (1968) Structure of a Mammalian Serine tRNA. Nature, 219, 1363–1365.

13. Kruppa, J. and Zachau, H.G. (1972) Multiplicity of serine-specific transfer RNAs of brewer’s and baker’s yeast. Biochimica et Biophysica Acta (BBA) - Nucleic Acids and Protein Synthesis, 277, 499–512.

14. Arango, D., Sturgill, D., Alhusaini, N., Dillman, A.A., Sweet, T.J., Hanson, G., Hosogane, M., Sinclair, W.R., Nanan, K.K., Mandler, M.D., et al. (2018) Acetylation of Cytidine in mRNA Promotes Translation Efficiency. Cell, 175, 1872–1886.e24.

15. Kudrin, P., Singh, A., Meierhofer, D., Kuśnierczyk, A. and Ørom, U.A.V. (2024) N4-acetylcytidine (ac4C) promotes mRNA localization to stress granules. EMBO Rep, 25, 1814–1834.

16. Liu, R., Wubulikasimu, Z., Cai, R., Meng, F., Cui, Q., Zhou, Y. and Li, Y. (2023) NAT10-mediated N4-acetylcytidine mRNA modification regulates self-renewal in human embryonic stem cells. Nucleic Acids Res, 51, 8514–8531.

17. Ito, S., Horikawa, S., Suzuki, T., Kawauchi, H., Tanaka, Y., Suzuki, T. and Suzuki, T. (2014) Human NAT10 Is an ATP-dependent RNA Acetyltransferase Responsible for *N*4-Acetylcytidine Formation in 18 S Ribosomal RNA (rRNA)*. Journal of Biological Chemistry, 289, 35724–35730.

18. Sharma, S., Langhendries, J.-L., Watzinger, P., Kötter, P., Entian, K.-D. and Lafontaine, D.L.J. (2015) Yeast Kre33 and human NAT10 are conserved 18S rRNA cytosine acetyltransferases that modify tRNAs assisted by the adaptor Tan1/THUMPD1. Nucleic Acids Res, 43, 2242–2258.

19. Ito, S., Akamatsu, Y., Noma, A., Kimura, S., Miyauchi, K., Ikeuchi, Y., Suzuki, T. and Suzuki, T. (2014) A Single Acetylation of 18 S rRNA Is Essential for Biogenesis of the Small Ribosomal Subunit in Saccharomyces cerevisiae. J Biol Chem, 289, 26201–26212.

20. Arango, D., Sturgill, D., Yang, R., Kanai, T., Bauer, P., Roy, J., Wang, Z., Hosogane, M., Schiffers, S. and Oberdoerffer, S. (2022) Direct epitranscriptomic regulation of mammalian translation initiation through N4-acetylcytidine. Mol Cell, 82, 2797–2814.e11.

21. Arango, D., Sturgill, D., Alhusaini, N., Dillman, A.A., Sweet, T.J., Hanson, G., Hosogane, M., Sinclair, W.R., Nanan, K.K., Mandler, M.D., et al. (2018) Acetylation of Cytidine in mRNA Promotes Translation Efficiency. Cell, 175, 1872–1886.e24.

22. Dang, Y., Li, J., Li, Y., Wang, Y., Zhao, Y., Zhao, N., Li, W., Zhang, H., Ye, C., Ma, H., et al. N-acetyltransferase 10 regulates alphavirus replication via N4-acetylcytidine (ac4C) modification of the lymphocyte antigen six family member E (LY6E) mRNA. J Virol, 98, e01350–23.

23. Hao, H., Liu, W., Miao, Y., Ma, L., Yu, B., Liu, L., Yang, C., Zhang, K., Chen, Z., Yang, J., et al. (2022) N4-acetylcytidine regulates the replication and pathogenicity of enterovirus 71. Nucleic Acids Research, 50, 9339–9354.

24. Yan, Q., Zhou, J., Wang, Z., Ding, X., Ma, X., Li, W., Jia, X., Gao, S.-J. and Lu, C. (2023) NAT10- dependent N4-acetylcytidine modification mediates PAN RNA stability, KSHV reactivation, and IFI16-related inflammasome activation. Nat Commun, 14, 6327.

25. Furman, D., Chang, J., Lartigue, L., Bolen, C.R., Haddad, F., Gaudilliere, B., Ganio, E.A., Fragiadakis, G.K., Spitzer, M.H., Douchet, I., et al. (2017) Expression of specific inflammasome gene modules stratifies older individuals into two extreme clinical and immunological states. Nat Med, 23, 174–184.

26. Duan, J., Zhang, Q., Hu, X., Lu, D., Yu, W. and Bai, H. (2019) N4-acetylcytidine is required for sustained NLRP3 inflammasome activation via HMGB1 pathway in microglia. Cellular Signalling, 58, 44–52.

27. Szymańska, E., Markuszewski, M.J., Markuszewski, M. and Kaliszan, R. (2010) Altered levels of nucleoside metabolite profiles in urogenital tract cancer measured by capillary electrophoresis. Journal of Pharmaceutical and Biomedical Analysis, 53, 1305–1312.

28. Li, H., Qin, Q., Shi, X., He, J. and Xu, G. (2019) Modified metabolites mapping by liquid chromatography-high resolution mass spectrometry using full scan/all ion fragmentation/neutral loss acquisition. Journal of Chromatography A, 1583, 80–87.

29. Xu, C., Zhang, J., Zhang, J. and Liu, B. (2021) SIRT7 is a deacetylase of N4-acetylcytidine on ribosomal RNA. GENOME INSTAB. DIS., 2, 253–260.

30. Oashi, Z., Murao, K., Yahagi, T., Von Minden, D.L., McCloskey, J.A. and Nishimura, S. (1972) Characterization of C + located in the first position of the anticodon of Escherichia coli tRNA Met as N 4 -acetylcytidine. Biochim Biophys Acta, 262, 209–213.

31. Ikeuchi, Y., Kitahara, K. and Suzuki, T. (2008) The RNA acetyltransferase driven by ATP hydrolysis synthesizes N4-acetylcytidine of tRNA anticodon. EMBO J, 27, 2194–2203.

32. Taniguchi, T., Miyauchi, K., Sakaguchi, Y., Yamashita, S., Soma, A., Tomita, K. and Suzuki, T. (2018) Acetate-dependent tRNA acetylation required for decoding fidelity in protein synthesis. Nat Chem Biol, 14, 1010–1020.

33. Orita, I., Futatsuishi, R., Adachi, K., Ohira, T., Kaneko, A., Minowa, K., Suzuki, M., Tamura, T., Nakamura, S., Imanaka, T., et al. (2019) Random mutagenesis of a hyperthermophilic archaeon identified tRNA modifications associated with cellular hyperthermotolerance. Nucleic Acids Res, 47, 1964–1976.

34. Sas-Chen, A., Thomas, J.M., Matzov, D., Taoka, M., Nance, K.D., Nir, R., Bryson, K.M., Shachar, R., Liman, G.L.S., Burkhart, B.W., et al. (2020) Dynamic RNA acetylation revealed by quantitative cross-evolutionary mapping. Nature, 583, 638–643.

35. Wolff, P., Villette, C., Zumsteg, J., Heintz, D., Antoine, L., Chane-Woon-Ming, B., Droogmans, L., Grosjean, H. and Westhof, E. (2020) Comparative patterns of modified nucleotides in individual tRNA species from a mesophilic and two thermophilic archaea. RNA, 26, 1957–1975.

36. Bruenger, E., Kowalak, J.A., Kuchino, Y., McCloskey, J.A., Mizushima, H., Stetter, K.O. and Crain, P.F. (1993) 5S rRNA modification in the hyperthermophilic archaea Sulfolobus solfataricus and Pyrodictium occultum. FASEB J, 7, 196–200.

37. Ohira, T. and Suzuki, T. (2024) Transfer RNA modifications and cellular thermotolerance. Molecular Cell, 84, 94–106.

38. Statkevičiūtė, R., Sadauskas, M., Rainytė, J., Kavaliauskaitė, K. and Meškys, R. (2022) Comparative Analysis of Mesophilic YqfB-Type Amidohydrolases. Biomolecules, 12, 1492.

39. Stanislauskienė, R., Laurynėnas, A., Rutkienė, R., Aučynaitė, A., Tauraitė, D., Meškienė, R., Urbelienė, N., Kaupinis, A., Valius, M., Kaliniene, L., et al. (2020) YqfB protein from Escherichia coli: an atypical amidohydrolase active towards N4-acylcytosine derivatives. Sci Rep, 10, 788.

40. Meng, C., Shi, X., Guo, W., Jian, X., Zhao, J., Wen, Y., Wang, R., Li, Y., Xu, S., Chen, H., et al. (2025) Structural analysis of ASCH domain-containing proteins and their implications for nucleotide processing. Structure, 33, 2095–2108.e5.

41. Iyer, L.M., Burroughs, A.M. and Aravind, L. (2006) The ASCH superfamily: novel domains with a fold related to the PUA domain and a potential role in RNA metabolism. Bioinformatics, 22, 257–263.

42. Jung, D.-J., Sung, H.-S., Goo, Y.-W., Lee, H.M., Park, O.K., Jung, S.-Y., Lim, J., Kim, H.-J., Lee, S.-K., Kim, T.S., et al. (2002) Novel transcription coactivator complex containing activating signal cointegrator 1. Mol Cell Biol, 22, 5203–5211.

43. Kim, H.-J., Yi, Ji-Young, Sung, Hee-Sook, Moore, David D., Jhun, Byung Hak, Lee, Young Chul and and Lee, J.W. (1999) Activating Signal Cointegrator 1, a Novel Transcription Coactivator of Nuclear Receptors, and Its Cytosolic Localization under Conditions of Serum Deprivation. Molecular and Cellular Biology, 19, 6323–6332.

44. Yoo, H.M., Kang, S.H., Kim, J.Y., Lee, J.E., Seong, M.W., Lee, S.W., Ka, S.H., Sou, Y.-S., Komatsu, M., Tanaka, K., et al. (2014) Modification of ASC1 by UFM1 Is Crucial for ERα Transactivation and Breast Cancer Development. Molecular Cell, 56, 261–274.

45. Hu, C., Chen, Z., Wang, G., Yang, H. and Ding, J. (2024) Biochemical and structural characterization of the DNA-binding properties of human TRIP4 ASCH domain reveals insights into its functional role. Structure, 32, 1208–1221.e4.

46. Kim, M., Park, S.H., Park, J.S., Kim, H.-J. and Han, B.W. (2019) Crystal Structure of Human EOLA1 Implies Its Possibility of RNA Binding. Molecules, 24, 3529.

47. Liu, Y., Liu, H., Chen, W., Yang, T. and Zhang, W. (2014) EOLA1 protects lipopolysaccharide induced IL-6 production and apoptosis by regulation of MT2A in human umbilical vein endothelial cells. Mol Cell Biochem, 395, 45–51.

48. Shi, X., Zhang, Y., Liu, N., Wang, R., Zhang, N., Cao, Y., Wang, D., Jin, Y., Meng, Q., Fan, S., et al. (2026) EOLA1, a novel mitochondria-localized protein critical for heart functions via regulating mitochondrial translation. 10.64898/2026.01.12.699056.

49. Kim, B.-N., Shin, M., Ha, S.C., Park, S.-Y., Seo, P.-W., Hofmann, A. and Kim, J.-S. (2017) Crystal structure of an ASCH protein from Zymomonas mobilis and its ribonuclease activity specific for single-stranded RNA. Sci Rep, 7, 12303.

50. Park, S.-Y., Park, J.-H. and Kim, J.-S. (2011) Cloning, expression, purification, crystallization and preliminary X-ray diffraction analysis of an ASCH domain-containing protein from Zymomonas mobilis ZM4. Acta Crystallogr Sect F Struct Biol Cryst Commun, 67, 310–312.

51. Laemmli, U.K. (1970) Cleavage of structural proteins during the assembly of the head of bacteriophage T4. Nature, 227, 680–685.

52. Blum, M., Andreeva, A., Florentino, L.C., Chuguransky, S.R., Grego, T., Hobbs, E., Pinto, B.L., Orr, A., Paysan-Lafosse, T., Ponamareva, I., et al. (2025) InterPro: the protein sequence classification resource in 2025. Nucleic Acids Res, 53, D444–D456.

53. Fu, L., Niu, B., Zhu, Z., Wu, S. and Li, W. (2012) CD-HIT: accelerated for clustering the next-generation sequencing data. Bioinformatics, 28, 3150–3152.

54. Wu, R., Ding, F., Wang, R., Shen, R., Zhang, X., Luo, S., Su, C., Wu, Z., Xie, Q., Berger, B., et al. (2022) High-resolution de novo structure prediction from primary sequence. 10.1101/2022.07.21.500999.

55. Zhang, Y. and Skolnick, J. (2005) TM-align: a protein structure alignment algorithm based on the TM-score. Nucleic Acids Res, 33, 2302–2309.

56. Talevich, E., Invergo, B.M., Cock, P.J. and Chapman, B.A. (2012) Bio.Phylo: A unified toolkit for processing, analyzing and visualizing phylogenetic trees in Biopython. BMC Bioinformatics, 13, 209.

57. Huerta-Cepas, J., Serra, F. and Bork, P. (2016) ETE 3: Reconstruction, Analysis, and Visualization of Phylogenomic Data. Mol Biol Evol, 33, 1635–1638.

58. Abramson, J., Adler, J., Dunger, J., Evans, R., Green, T., Pritzel, A., Ronneberger, O., Willmore, L., Ballard, A.J., Bambrick, J., et al. (2024) Accurate structure prediction of biomolecular interactions with AlphaFold 3. Nature, 630, 493–500.

59. Holm, L., Laiho, A., Törönen, P. and Salgado, M. (2023) DALI shines a light on remote homologs: One hundred discoveries. Protein Science, 32, e4519.

60. Letunic, I. and Bork, P. (2024) Interactive Tree of Life (iTOL) v6: recent updates to the phylogenetic tree display and annotation tool. Nucleic Acids Res, 52, W78–W82.

61. UniProt Consortium (2025) UniProt: the Universal Protein Knowledgebase in 2025. Nucleic Acids Res, 53, D609–D617.

62. Letunic, I. and Bork, P. (2026) SMART v10: three decades of the protein domain annotation resource. Nucleic Acids Res, 54, D499–D503.

63. Zimmermann, L., Stephens, A., Nam, S.-Z., Rau, D., Kübler, J., Lozajic, M., Gabler, F., Söding, J., Lupas, A.N. and Alva, V. (2018) A Completely Reimplemented MPI Bioinformatics Toolkit with a New HHpred Server at its Core. Journal of Molecular Biology, 430, 2237–2243.

64. Jayaraman, V., Toledo-Patiño, S., Noda-García, L. and Laurino, P. (2022) Mechanisms of protein evolution. Protein Science, 31, e4362.

65. Toledo-Patiño, S., Pascarelli, S., Uechi, G. and Laurino, P. Insertions and deletions mediated functional divergence of Rossmann fold enzymes. Proc Natl Acad Sci U S A, 119, e2207965119.

66. Chan, P., Curtis, R.A. and Warwicker, J. (2013) Soluble expression of proteins correlates with a lack of positively-charged surface. Sci Rep, 3, 3333.

67. Alayyoubi, M., Guo, H., Dey, S., Golnazarian, T., Brooks, G.A., Rong, A., Miller, J.F. and Ghosh, P. (2013) Structure of the essential diversity-generating retroelement protein bAvd and its functionally important interaction with reverse transcriptase. Structure, 21, 266–276.

68. Du, H., Zhao, Y., He, J., Zhang, Y., Xi, H., Liu, M., Ma, J. and Wu, L. (2016) YTHDF2 destabilizes m6A-containing RNA through direct recruitment of the CCR4–NOT deadenylase complex. Nat Commun, 7, 12626.

69. Zhang, Z., Theler, D., Kaminska, K.H., Hiller, M., de la Grange, P., Pudimat, R., Rafalska, I., Heinrich, B., Bujnicki, J.M., Allain, F.H.-T., et al. (2010) The YTH Domain Is a Novel RNA Binding Domain. J Biol Chem, 285, 14701–14710.

70. Wang, S., Xie, H., Mao, F., Wang, H., Wang, S., Chen, Z., Zhang, Y., Xu, Z., Xing, J., Cui, Z., et al. (2022) N4-acetyldeoxycytosine DNA modification marks euchromatin regions in Arabidopsis thaliana. Genome Biology, 23, 5.

71. Anantharaman, V., Zhang, D. and Aravind, L. (2010) OST-HTH: a novel predicted RNA-binding domain. Biol Direct, 5, 13.

72. Kubíková, J., Reinig, R., Salgania, H.K. and Jeske, M. (2021) LOTUS-domain proteins - developmental effectors from a molecular perspective. Biological Chemistry, 402, 7–23.

73. Chujo, T. and Tomizawa, K. (2025) Mitochondrial tRNA modifications: functions, diseases caused by their loss, and treatment strategies. RNA, 31, 382–394.

74. Preitakaitė, V., Barasa, P., Aučynaitė, A., Plakys, G., Koplūnaitė, M., Zubavičiūtė, S. and Meškys, R. (2023) Bacterial amidohydrolases and modified 5-fluorocytidine compounds: Novel enzyme-prodrug pairs. PLoS One, 18, e0294696.

