## Supplementary material for "ASCH Domain-Containing Proteins Act as tRNA *N*^4^-acetylcytidine Erasers": Statkeviciute_et_al_supplementary_data.pdf

**Supplementary Table S1:** list of oligonucleotide primers used for gene amplification.

| Primer name | Sequence (5′→3′) | Purpose |
|---|---|---|
| Cun_trunc_F | AGAAGGAGATATAACTATGAAAAACGACATCACATTCTAC | CunASCH N-terminus truncation |
| CunYqfB_R | GTGGTGGTGATGGTGATGGCCTTCATCAATCTTTTTAAAAGC | |
| Afu1ASCH_F | AGAAGGAGATATAACTATGAGAGAGCCCTTTGCATCCATGATAG | Amplification of the target Afu1ASCH gene for LIC cloning into pLATE31 |
| Afu1ASCH_R | GTGGTGGTGATGGTGATGGCCGTCAATCTTAACCCA | |
| Afu2ASCH_F | AGAAGGAGATATAACTATGGAGAGGATTAACTTT | Amplification of the target Afu2ASCH gene for LIC cloning into pLATE31 |
| Afu2ASCH_R | GTGGTGGTGATGGTGATGGCCTTTCACGACCTCGAA | |
| Afu3ASCH_F | AGAAGGAGATATAACTATGAAGCACCTTGAGTTT | Amplification of the target Afu3ASCH gene for LIC cloning into pLATE31 |
| Afu3ASCH_R | GTGGTGGTGATGGTGATGGCCAATCAGCCCCCTCTT | |
| TthASCH_F | AGAAGGAGATATAACTATGGAAAGGCCCAAG | Amplification of the target TthASCH gene for LIC cloning into pLATE31 |
| TthASCH_R | GTGGTGGTGATGGTGATGGCCCCACCGTACCTCGGA | |
| Afu3ASCH_F | AGAAGGAGATATAACTATGAAGCACCTTGAGTTT | |

| | | |
|---|---|---|
| Afu3_trunc_R | GTGGTGGTGATGGTGATGGCCAGGCTCAAAATCGAAGTGTATGAC | Amplification of the stand-alone ASCH domain of Afu3ASCH for LIC cloning into pLATE31 |
| Pku4ASCH_F | AGAAGGAGATATAACTATGAAAAACCTGAAATTCGATGGACG | Amplification of the stand-alone ASCH domain of Pku4ASCH for LIC cloning into pLATE31 |
| Pku4_trunc_R | GTGGTGGTGATGGTGATGGCCTTTCACCATCTCAAATTCAACGATG | |

**Supplementary Table S2:** list of DNA oligonucleotide primers used for site-directed mutagenesis.

| Primer name | Sequence (5′→3′) | Template |
|---|---|---|
| Tth_Y14A_F | CCGCAGCCAGCCTCATCGTGGACGGGA | pLATE31-TthASCH |
| Tth_Y14A_R | CTGCGGGTTCCCGGACGATGAGGCCGA | |
| Tth_Y14F_F | CCTTCGCCAGCCTCATCGTGGACGGGA | |
| Tth_Y14F_R | CGAAGGGTTCCCGGACGATGAGGCCGA | |
| Tth_K23A_F | AGGGCGGTCTGGGAGATCCGCAGG | |
| Tth_K23A_R | GACCCGCCTCCCGTCCACGATGA | |
| Tth_K23L_F | AGGCTGGTCTGGGAGATCCGCAGG | |
| Tth_K23L_R | GACCAGCCTCCCGTCCACGATGA | |
| Tth_E26A_F | TGGGCGATCCGCAGGCGCAAAACCCGC | |
| Tth_E26A_R | GATCGCCCAGACCTTCCTCCCGTCCACGAT | |
| Tth_E26D_F | TGGGACATCCGCAGGCGCAAAACCCGC | |
| Tth_E26D_R | GATGTCCCAGACCTTCCTCCCGTCCACGAT | |
| Tth_E26S_F | TGGTCCATCCGCAGGCGCAAAACCCGC | |
| Tth_E26S_R | GATGGACCAGACCTTCCTCCCGTCCACGAT | |
| Tth_E26T_F | TGGACCATCCGCAGGCGCAAAACCCGC | |

| | | |
|---|---|---|
| Tth_E26T_R | TGGTCCAGACCTTCCTCCCGTCCACGATGA | |
| Tth_R28A_F | ATCGCCAGGCGCAAAACCCGCCACCG | |
| Tth_R28A_R | CCTGGCGATCTCCCAGACCTTCCTCCCGT | |
| Tth_R28L_F | ATCCTGAGGCGCAAAACCCGCCACCG | |
| Tth_R28L_R | CCTCAGGATCTCCCAGACCTTCCTCCCGT | |
| Tth_Y81A_F | CCGCTGCCAAGGACGAGCCCCTCTACGC | |
| Tth_Y81A_R | CAGCGGCCCGGAGGAAGGCCTCCTCC | |
| Tth_Y81F_F | CTTCGCCAAGGACGAGCCCCTCTACGCC | |
| Tth_Y81F_R | GAAGGCCCGGAGGAAGGCCTCCTCCGC | |
| Tth_Y81L_F | CCTCGCCAAGGACGAGCCCCTCTACGCC | |
| Tth_Y81L_R | GCGAGGGCCCGGAGGAAGGCCTCCTCC | |
| EOLA1_K21A_F | GGCATCGCGACTGTTGAAACTCGTTGG | pET29-EOLA1 |
| EOLA1_K21A_R | GTTCAAAACAAATCCCGCGTATGGTTGGC | |
| EOLA1_E24A_F | GGCATCAAAACTGTTGCGACTCGTTGGC | |
| EOLA1_E24A_R | GTTCAAAACAAATCCCGCGTATGGTTGGC | |
| EOLA1_E24D_F | GGCATCAAAACTGTTGATACTCGTTGGC | |
| EOLA1_E24D_R | GTTCAAAACAAATCCCGCGTATGGTTGGC | |
| EOLA1_E24S_F | GGCATCAAAACTGTTTCCACTCGTTGGC | |
| EOLA1_E24S_R | GTTCAAAACAAATCCCGCGTATGGTTGGC | |
| EOLA1_R26A_F | GTTGAAACTGCGTGGCGGCCACTGCTG | |
| EOLA1_R26A_R | AGTTTTGATGCCGTTCAAAACAAATCCCGCGTATG | |
| EOLA1_E24K_F | GGCATCAAAACTGTTAAAACTCGTTGG | |
| EOLA1_E24K_R | GTTCAAAACAAATCCCGCGTATGGTTG | |
| EOLA1_T25M_F | GGCATCAAAACTGTTGAAATGCGTTGG | |
| EOLA1_T25M_R | GTTCAAAACAAATCCCGCGTATGGTTG | |

| | | |
|---|---|---|
| EOLA1_R26C_F | GGCATCAAAACTGTTGAAACTTGCTGGCG | |
| EOLA1_R26C_R | GTTCAAAACAAATCCCGCGTATGGTTGGCG | |
| EOLA1_R28H_F | GTTGAAACTCGTTGGCATCCACTGCTG | |
| EOLA1_R28H_R | AGTTTTGATGCCGTTCAAAACAAATCCC | |
| TRIP4ASCH_G16W_F | CTGGTGCGTTGGATCAAACGGGTTGAAG | pET21-TRIP4ASCH_NHis |
| TRIP4ASCH_K18A_F | CTGGTGCGTGGAATCGCACGGGTTG | |
| TRIP4ASCH_G16_K18_R | CAGACTCGCCCACGGCTGATG | |
| TRIP4ASCH_E21A_F | GTTGCGGGTCGCTCATGGTATACCC | |
| TRIP4ASCH_R23A_F | GTTGAAGGTGCGTCATGGTATACCCCG | |
| TRIP4ASCH_S24F_F | GTTGAAGGTCGCTTTTGGTATACCCCGCATC | |
| TRIP4ASCH_E21_R23_S24_R | CCGTTTGATTCCACGCACCAGCAG | |

**Supplementary Table S3:** list of oligonucleotide primers used for fusion protein generation.

| Primer name | Sequence (5′→3′) | Purpose |
|---|---|---|
| sfGFP_p51_F | GGTGATGATGATGACAAGCGTAAAGGCGAAGAGCT | Used in the first PCR round to amplify sfGFP;<br>Used in the second PCR round to amplify the final gene fusion product |
| GFP_WQ_Afu3_R | GCCTGTTGCAATTCCCAACCTTTGTACAGTTCATCCATACCATGCGT | Used in the first round of PCR to amplify sfGFP with WELQut recognition sequence and Afu3- or Pku4ASCH complementary flanks |
| GFP_WQ_Pku4_R | AACTTTTTGCAATTCCCAACCTTTGTACAGTTCATCCATACCATGCGT | |
| Afu3_WQ_GFP_F | TACAAAGGTTGGGAATTGCAACAGGCCGTCAATCCTCACAAGCT | |

| | | |
|---|---|---|
| Pku4_WQ_GFP_F | TACAAAGGTTGGGAATTGCAAAAAGTTCTGGATAAGCCTATTCTGT | Used in the first round of PCR to amplify Afu3- or Pku4ASCH with WELQut recognition sequence and sfGFP complementary flanks |
| Afu3_p51_R | GGAGATGGGAAGTCATTAAATCAGCCCCCTCTTCAC | Used in the first PCR round to amplify Afu3- or Pku4ASCH HTH domain;<br>Used in the second PCR round to amplify the final fusion product |
| Pku4_p51_R | GGAGATGGGAAGTCATTAAATTTTCGGTTCCATAATACCGCGCTT | |

**Supplementary Figure S4:** list of expression constructs used in this study. Construct sequences are available on Benchling.com via the provided links.

| Construct | Protein UniProt ID | Comments | Reference |
|---|---|---|---|
| pLATE11 | — | — | (1) |
| pET21a-YqfB | P67603 | C-terminal 6×His-tag<br>Amp$^R$ | (2) |
| pLATE31-BagASCH | UPI0015605FBC<br>(UniParc) | | (3) |
| pLATE31-CunASCH | A0AAC8VS57 | | |
| pLATE31-KpnASCH | A6TDQ4 | | |
| pLATE31-SloASCH | A3QD13 | | |
| pLATE31-BmyASCH | A0A084J0D7 | | This study |
| pLATE31-TthASCH | Q5SM30 | | |
| pLATE31-Afu1ASCH | O29197 | | |
| pLATE31-Afu2ASCH | O28749 | | |

| | | | |
|---|---|---|---|
| pLATE31-Afu3ASCH | O28463 | | |
| pET21-ZmASCH<br><br>construct sequence is available here | A0A0H3G0N3 | | (4) |
| pET21-TRIP4ASCH_NHis<br><br>construct sequence is available here | Q15650<br><br>(residues 435-581) | N-terminal 6×His-tag<br><br>$Amp^R$ | (5) |
| pET29b-Pku1ASCH<br><br>construct sequence is available here | A0A127B9I2 | C-terminal 6×His-tag<br><br>$Kan^R$ | This study |
| pET29b-Pku2ASCH<br><br>construct sequence is available here | A0A127BB00 | | |
| pET29b-Pku3ASCH<br><br>construct sequence is available here | A0A127BBJ2 | | |
| pET29b-Pku4ASCH<br><br>construct sequence is available here | A0A127B833 | | |
| pET29b-Pku5ASCH<br><br>construct sequence is available here | A0A127BC91 | | |
| pET29b-Pku6ASCH<br><br>construct sequence is available here | A0A127B787 | | |
| pET29b-EOLA1 isoform 1<br><br>construct sequence is available here | Q8TE69 | | |

<table>
<tr><td>pLATE51-GFP-Afu3_HTH<br><br>construct sequence is available here</td><td rowspan="6">—</td><td rowspan="2">N-terminal 6×His-tag<br><br>Amp$^R$</td><td rowspan="6">This study</td></tr>
<tr><td>pLATE51-GFP-Pku4_HTH<br><br>construct sequence is available here</td></tr>
<tr><td>pLATE31-YqfB-Afu3_HTH<br><br>construct sequence is available here</td><td rowspan="2">C-terminal 6×His-tag<br><br>Amp$^R$</td></tr>
<tr><td>pLATE31-YqfB-Pku4_HTH<br><br>construct sequence is available here</td></tr>
<tr><td>pLATE31-Afu3_tr<br><br>construct sequence is available here</td><td rowspan="2">C-terminal 6×His-tag<br><br>Amp$^R$</td></tr>
<tr><td>pLATE-Pku4_tr<br><br>construct sequence in available here</td></tr>
</table>

**Supplementary Table S5:** list of nucleic acid substrates used in this study.

<table>
<tr><th>Substrate name</th><th>Sequence (5′→3′)</th><th>Reference</th></tr>
<tr><td>ssRNA_17mer (Figure 6e)</td><td>CCCGACAACAGGCCCCC</td><td rowspan="3">(4)</td></tr>
<tr><td>ssRNA_30mer (Figure 6e)</td><td>AUCAGCUCGUCACAACAUUACUUCAUCAAC</td></tr>
<tr><td>ssDNA_30mer (Figure 6e)</td><td>ATCAGCTCGTCACAACATTACTTCATCAAC</td></tr>
<tr><td>ssDNA_24mer_F (Figure 4c, d)</td><td>FAM-TGATCATACCTCTGATCATACCTC</td><td rowspan="2">(5)</td></tr>
<tr><td>ssDNA_24mer_R (Figure 4d)</td><td>GAGGTATGATCAGAGGTATGATCA</td></tr>
</table>

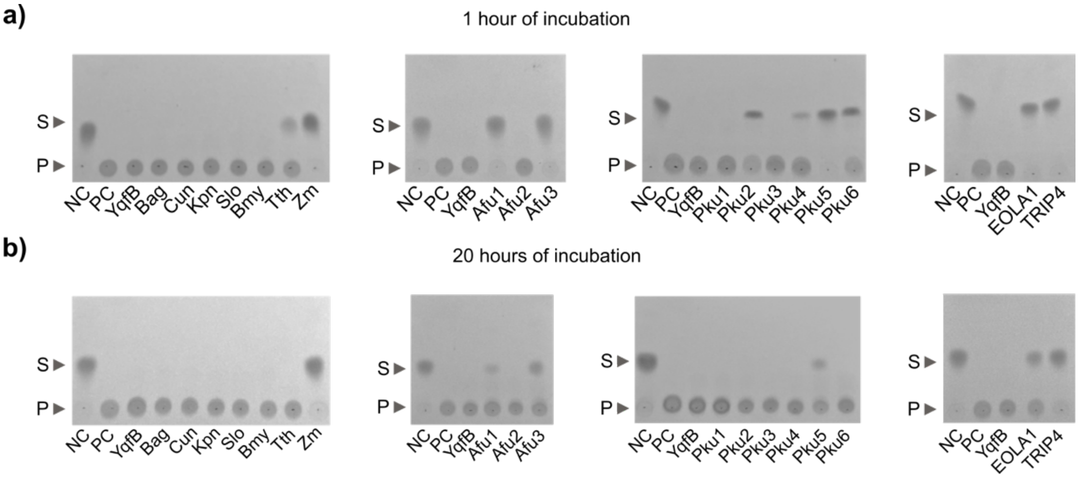


**Supplementary Figure S1:** TLC analysis of the amidohydrolytic activity of ASCH domain-containing proteins toward ac4C after 1 hour **(a)** and 20 hours **(b)** of incubation. S – reaction substrate, P – product.

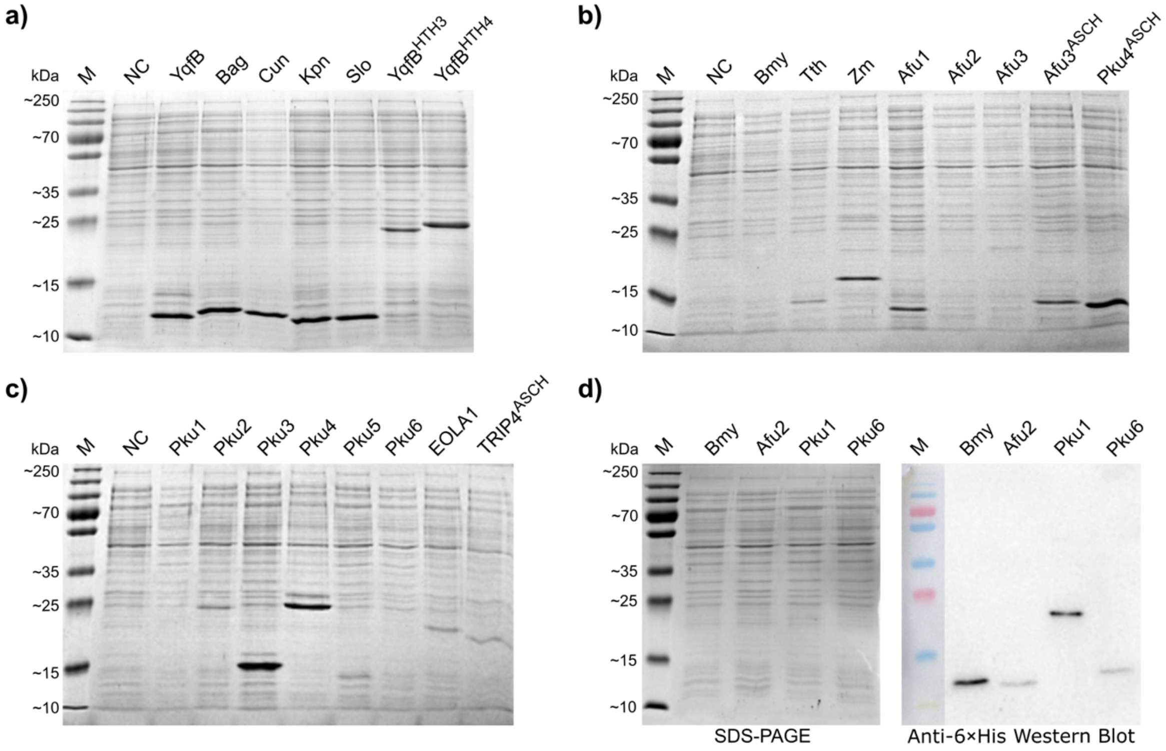


**Supplementary Figure S2: Analysis of soluble ASCH protein fractions in *E. coli* lysates. a-c)** SDS–PAGE analysis of soluble protein fractions obtained after 4 h of induction in *E. coli* KRX strain. **d)** Western blot analysis of the same soluble protein fractions used for SDS–PAGE, performed for constructs in which no visible protein bands were detected, confirming the presence of low amounts of soluble protein. M – PageRuler Prestained Protein Ladder, NC – negative control (lysate of *E. coli* KRX transformed with the empty pLATE11 vector).

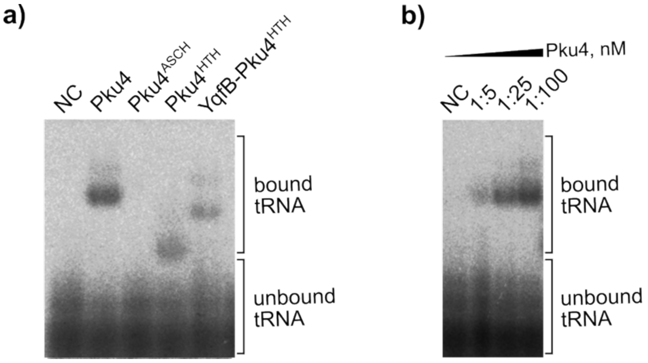


**Supplementary Figure S3: a)** tRNA binding analysis of Pku4ASCH-derived variants: Pku4 (full-length protein containing both ASCH and HTH domains), Pku4$^{ASCH}$ (stand-alone ASCH domain), Pku4$^{HTH}$ (standalone HTH domain), and YqfB-Pku4$^{HTH}$ fusion protein. **b)** EMSA analysis showing increased tRNA–protein complex formation with increasing full-length Pku4ASCH protein concentration (tRNA:protein molar ratios of 1:5, 1:25, and 1:100).

**References**

1. Jamontas,R., Laurynėnas,A., Povilaitytė,D., Meškys,R. and Aučynaitė,A. (2024) RudS: bacterial desulfidase responsible for tRNA 4-thiouridine de-modification. *Nucleic Acids Res.*, **52**, 10543–10562.

2. Stanislauskienė,R., Laurynėnas,A., Rutkienė,R., Aučynaitė,A., Tauraitė,D., Meškienė,R., Urbelienė,N., Kaupinis,A., Valius,M., Kaliniene,L., *et al.* (2020) YqfB protein from Escherichia coli: an atypical amidohydrolase active towards N4-acylcytosine derivatives. *Sci. Rep.*, **10**, 788.

3. Statkevičiūtė,R., Sadauskas,M., Rainytė,J., Kavaliauskaitė,K. and Meškys,R. (2022) Comparative Analysis of Mesophilic YqfB-Type Amidohydrolases. *Biomolecules*, **12**, 1492.

4. Kim,B.-N., Shin,M., Ha,S.C., Park,S.-Y., Seo,P.-W., Hofmann,A. and Kim,J.-S. (2017) Crystal structure of an ASCH protein from Zymomonas mobilis and its ribonuclease activity specific for single-stranded RNA. *Sci. Rep.*, **7**, 12303.

5. Hu,C., Chen,Z., Wang,G., Yang,H. and Ding,J. (2024) Biochemical and structural characterization of the DNA-binding properties of human TRIP4 ASCH domain reveals insights into its functional role. *Struct. Lond. Engl. 1993*, **32**, 1208-1221.e4.
